# Arctic humpback whales respond to nutritional opportunities before migration

**DOI:** 10.1101/2022.10.13.511409

**Authors:** Lisa Elena Kettemer, Theresia Ramm, Fredrik Broms, Martin Biuw, Marie-Anne Blanchet, Sophie Bourgeon, Paul Dubourg, Anna C. J. Ellendersen, Mathilde Horaud, Joanna Kershaw, Patrick J. O. Miller, Nils Øien, Logan J. Pallin, Audun H. Rikardsen

**Author notes:** These authors contributed equally to this work (Lisa Elena Kettemer, Theresia Ramm).

## Abstract

Rapid climate change in Arctic and Subarctic ecosystems is altering the spatio-temporal dynamics and abundance of resources. Whether highly mobile predators can respond and match their movements to changed resource peaks remains largely unclear. In the last decade, humpback whales (*Megaptera novaeangliae*) established a new foraging site in fjords of northern Norway during the winter, outside of their presumed foraging season. We used photographic matching to show that whales first sighted during fall in the Barents Sea foraged in northern Norway from late October to February, staying for up to three months and showing high inter-annual return rates (up to 82%). The number of identified whales increased steadily from 2010 to 2016. Genetic sexing and hormone profiling in both areas suggest higher proportions of pregnancy and a female bias in Norwegian waters. This indicates that the new site may be particularly important for pregnant females, likely to improve body condition before migration. Our results suggest that baleen whales can respond to nutritional opportunities along their migration pathways, in some cases by extending their feeding season. This supports the idea that migrating marine mammals can access novel prey resources as part of their response to environmental changes.

## Introduction

The Arctic is warming rapidly with climate change. Changes in the physical environment cascade through the food web, causing shifts in the timing and spatial occurrence of prey aggregations important to arctic predators (Descamps et al., 2017; Meredith et al., 2019). Seasonal visitors to the area, such as migratory land mammals, birds, and marine mammals, rely heavily on predictable high seasonal productivity to fuel their year-round energy demands. Some of these migratory species have already adapted their spatio-temporal distribution and movement patterns in response to environmental change (Davidson et al., 2020). However, how well migrants can respond to such changes is unclear due to a limited understanding of the behavioral plasticity underlying migratory decisions and how these organisms can assess and adapt to complex dynamics in their environment. Most baleen whales show strong culturally transmitted philopatry to foraging and breeding grounds (Barendse et al., 2013) and likely base their movements on memory of past resource distributions (Abrahms et al., 2019). These adaptations enable migrants to match the timing of their movements to seasonal resource phenology. Some species of baleen whales have changed the timing of their migrations (Avila et al., 2020; Ramp et al., 2015) and distribution during the foraging season, re-populating historical foraging grounds decades after near-extirpation from overexploitation (Calderan et al., 2020; Keen et al., 2021).

In the last decade, humpback whales (*Megaptera novaeangliae*) have established a novel foraging site following increased occurrence of Norwegian spring-spawning (NSS) herring that overwinter in fjords of northern Norway during winter (Jourdain & Vongraven, 2017; Kettemer et al., 2022). Northeast Atlantic humpback whales forage throughout the Norwegian and Barents Seas during summer and autumn (Christensen et al., 1992; Hamilton et al., 2021; Leonard & Øien, 2020) and migrate to breeding grounds in the West Indies (Stevick et al., 2018) and Cape Verde Islands (Wenzel et al., 2020), where most of them are observed in March -April and April -May, respectively. During the era of commercial whaling in the Northeast Atlantic (1881 -1904), humpback whales were caught off northern Norway in areas occupied by herring during the winter (Christensen et al., 1992; Ingebrigtsen, 1929; Kramvig et al., 2016), but no substantial numbers of whales have been observed in the area since then. This novel or re-established foraging site appears to represent a continuation of the main foraging season or a feeding stopover on the southward migration, based on the timing and location about 1,000 kilometers southeast of the Barents Sea, but no formal analysis of the connectivity between the two areas has been conducted.

In this study, we aimed to assess the importance of the Norwegian fjords in providing continued foraging opportunities to various demographic groups of Northeast Atlantic humpback whales. To this end, we used photographic matching to (1) investigate the connectivity between the Barents Sea and the Norwegian fjords, and (2) to assess the return rate of individual whales that foraged in the fjords both within and between seasons. Finally, we used biopsy samples to (3) assess the sex ratio and pregnancy rate in both areas and throughout the season.

## Materials and Methods

### Study site and data collection

We collected photo-identification data and biopsies in several northern Norwegian fjords and waters surrounding the Svalbard Archipelago (Figure 1). The North Norwegian Humpback Whale Catalogue (NNHWC) was established in 2010 when humpback whales started aggregating in northern Norwegian fjords during the winter. The study sites included waters around Andøya (2010 to 2012), Kvaløya (2012 to 2017), and Kvænangen (2017 to 2019) (Figure 1). Photographic sampling was conducted using small vessels and was dictated by weather and light conditions. During the polar night (December-January), sampling was usually restricted to a few hours around midday. However, on some sampling trips, a flash system allowed sampling to continue in low light conditions. The sampling effort differed between years and study sites (Table 1). The public and other research organizations also submitted pictures, and an interactive online web portal for the submission of fluke photographs was established in 2015 (hvalid.no) and active until 2017.

**Figure 1.**
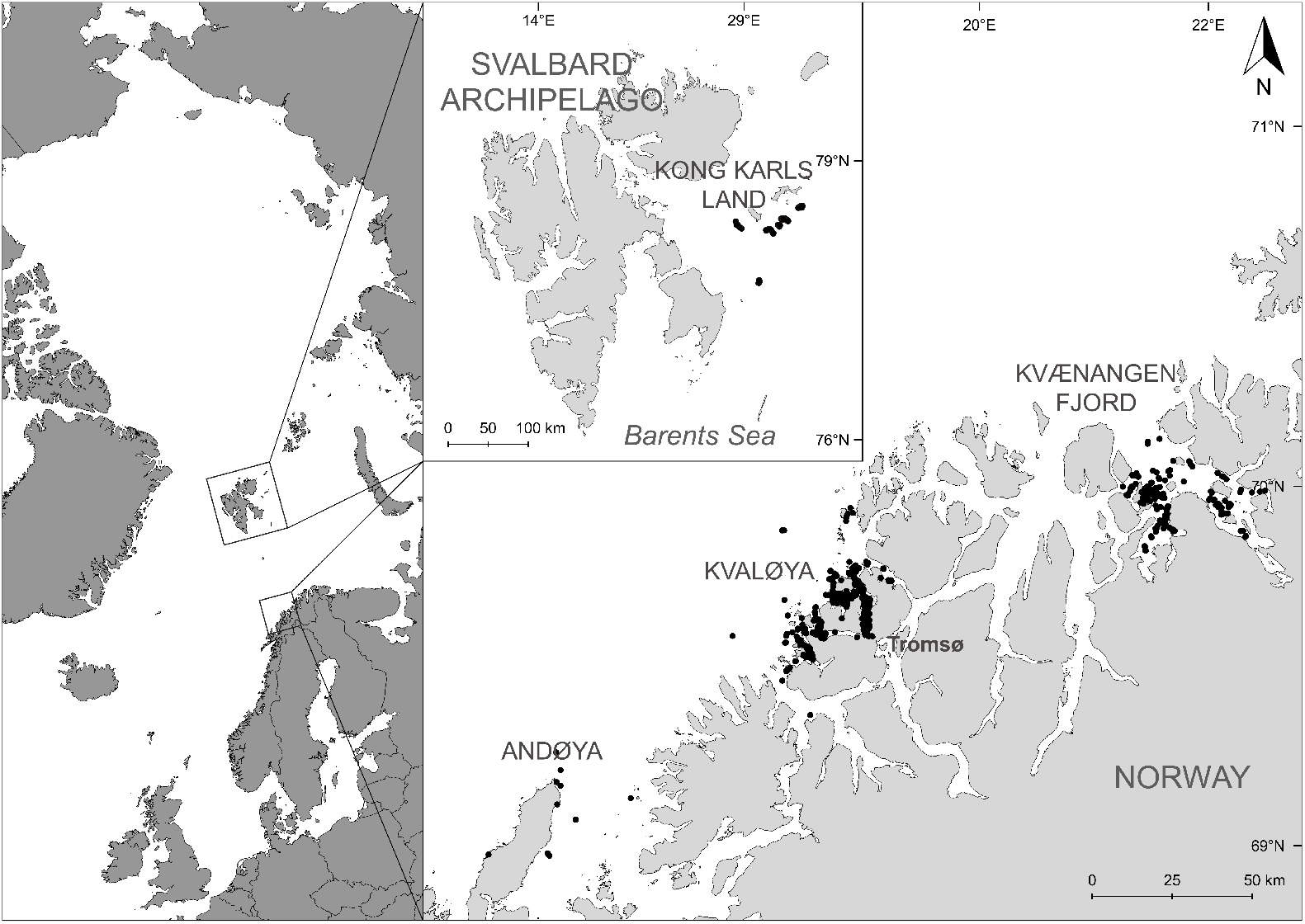
Map of sighted individuals with available GPS location. Left panel shows the Svalbard Archipelago with black dots close to Kong Karls Land representing exact GPS points of photographic records of humpback whales (*Megaptera novaeangliae*). Pictures without exact GPS locations are not included in the figure. The inset shows the three main locations (Andøya, Kvaløya, and Kvænangen fjord) of the northern Norwegian foraging area with exact humpback whale locations.

**Table 1.**
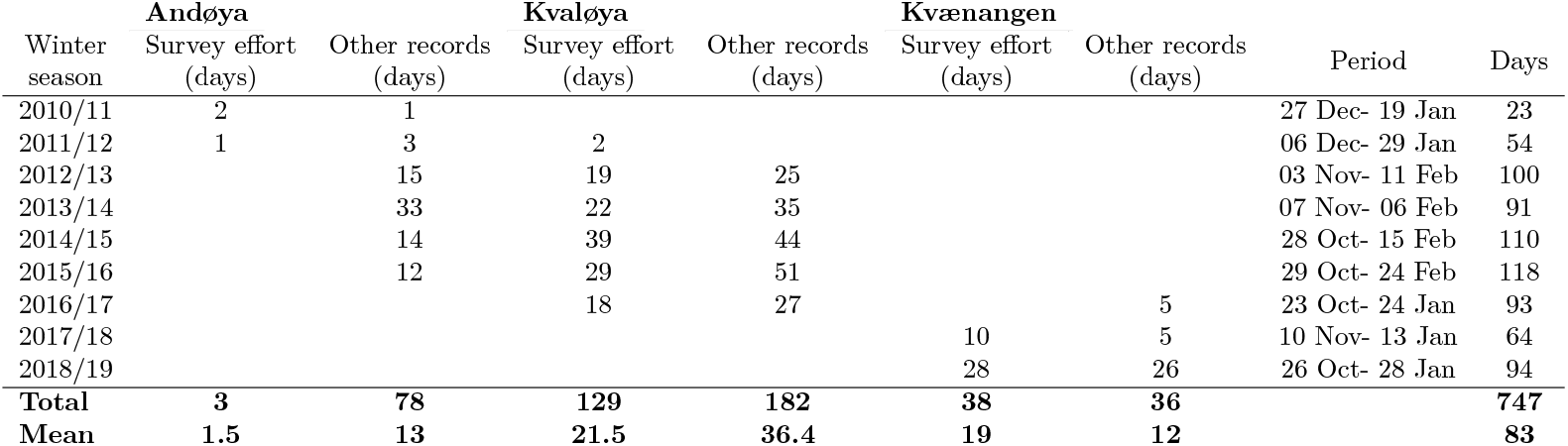
Table of effort-based sampling and non-effort-based data collection for each location within the northern Norwegian fjords (2010/11 to 2018/19). Sampling was conducted by the UiT and the founder of the NNHWC (effort-based). Other records (non-effort-based) represent days in which various contributors submitted fluke identification photographs. The period depicts the first and last humpback whale fluke capture in a season, with days indicating the duration between them, indicative of season duration.

From the 3rd to the 11th of September 2018, a research cruise was conducted in cooperation between the Institute of Marine Research (IMR Bergen, Norway) and the University of Tromsø (UiT, Tromsø, Norway), surveying the northern Barents Sea, east of the Svalbard archipelago, close to the island group of Kong Karls Land (Figure 1). We chose the timing and area based on information on humpback whale occurrence from prior annual joint Norwegian/Russian ecosystem surveys in the Barents Sea and adjacent waters (IMR, Norway/PINRO, Russia). When humpback whales were sighted, a small boat was launched to allow closer approaches. We took fluke photographs from both the small boat and the larger research vessel using DSLR cameras. In addition to this cruise, photographs from incidental humpback whale encounters around Svalbard, and the Barents Sea were submitted by various contributors (2012 to 2019).

We took biopsies from either the fluke or flank of each individual from small open boats (20-26ft) using an airgun (ARTS launching system, LKARTS-Norway) to deploy a floating arrow with a 4 or 6 cm long sterile stainless steel biopsy tip (CetaDart, DK). Depending on the shooting distance, usually about 4-20 meters, the shooting pressure was between 6-10 bars.

Sampling procedures were approved by the Norwegian Food Safety Authorities (Mattilsynet), under permits FOTS-ID 14135 and FOTS-ID 8165. We collected skin (N = 169) and blubber samples (N = 112) from humpback whales between 2011 -2019 in the Troms area of northern Norway, and during September 2018 in the northern Barents Sea. We stored samples at -20°C in either tin foil or glass vials (blubber) or 96% EtOH (skin).

### Photo-identification

We identified individual humpback whales using the unique pigmentation pattern on their ventral flukes (Katona & Whitehead, 1981). We created sighting histories from re-identifications of such photo-identified whales. Intervals between an individual’s first and last sightings within a season indicate the minimum length of stay during the season. We calculated the annual return rate, a measure of site fidelity on a population level, as the number of photographically recaptured individuals in a given year divided by the total number of individuals sighted in that year (Clapham et al., 1993).

Individual sighting histories for this study relied on 3677 sightings of 1169 unique humpback whales taken between 2010 and 2019, in the NNHWC. The catalog covers a latitudinal range from 67º to 80º N. It contains sighting records of individual humpback whales throughout the year, with summer and autumn sightings mainly focused in the Barents Sea and winter sightings in northern Norway (Figure 1). In northern Norway, we collected fluke photographs of 866 individual humpback whales, 856 of these during the winter. We collected most (54.7%) photographs during dedicated sampling conducted between October and February; others contributed the remaining pictures including all summer sightings (1%).

Over nine years of study, we conducted 170 days of dedicated photo-identification survey effort, with considerably less effort during the first two winters (Table 1). The average annual sampling effort across all winter seasons was 17 days (± 13.1) and 23.6 days (± 9.4), excluding the first two seasons. We identified 342 individual whales in the Barents Sea, with most identification photographs (95%) obtained during a research cruise in September 2018. Other collaborators submitted fluke photographs from incidental humpback whale encounters between 2012 and 2019.

### Sex determination

We determined the sex of individuals using skin samples (Bérubé & Palsbøll, 1996). We used the odontocete oligonucleotide primer set, ZFYX0582F, ZFY0767R and ZFX0923R, which showed clear bands on the gel electrophoresis. As a control, samples from four killer whales (*Orcinus orca*) of known sex (two males and two females) were used in every PCR reaction. After initial testing, primer concentrations were optimized to 1 µl of 10 µM for the Y primer-set (ZFYX0582F/ZFY0767R) and 0.5 µl of 5 µM for the X primer-set (ZFYX0582F/ZFX0923R).

### Relatedness

To estimate the within-season recapture rate in our dataset, we conducted a relatedness analysis on a subset of the samples for which genetic sequences were available (N= 107). We used NGSrelate v2 (Hanghøj et al., 2019) to calculate the coefficients of relatedness, based on genotype likelihoods calculated with ANGSD v.v0.935-53-gf475f10 (Korneliussen et al., 2014). See S2 Text in supporting information for more details.

### Progesterone concentrations and pregnancy status

We used progesterone concentrations as a proxy for pregnancy status and extracted the progesterone from blubber samples as described in (Kershaw et al., 2020; Pallin et al., 2018), with minor adjustments to the method. We measured progesterone in 82 female blubber samples and 19 male control samples. We excluded blubber samples taken from flukes since they usually do not contain enough blubber to conduct the analysis and may have different fat and hormone profiles leading to potential misclassifications. We thawed frozen blubber samples at room temperature and homogenized them seven times for 40 sec at 5000 rpm, using a Precellys 24 tissue homogenizer, and cooled samples on ice between intervals. We rinsed the resulting homogenates using a series of solvent washes, removed tissue debris, collected the supernatants, and evaporated the samples using pressurized air. We stopped evaporation when a thin solvent coating was left to prevent complete desiccation and potential oxidation of the residue. A subset of samples was homogenized manually before the same tissue debris removal, and solvent washes and extracts were dried down under nitrogen.

We quantified progesterone concentrations using two commercially available progesterone enzyme immunoassays (EIA; Enzo Life Sciences, kit ADI-900-011, and ELISA; DRG International Inc. EIA-1561), see S1 Text and S3 Table in supporting information for more details on the difference between the two methods. We re-suspended the dried hormone extract in 1ml phosphate buffered saline (pH 7.5) containing 1% bovine serum albumin, vortexed and then kept samples at -20°C. The EIA and ELISA kits we used have 100% reactivity with progesterone; the detection limit is between 15 -500 pg ml^1^ and 0 -40 ng ml^1^, respectively, based on the standard curves. We added two additional standard dilutions to lower the detection limit of the EIA standard curve to 3.81 pg ml^1^. We ran samples blind and in duplicate and re-ran samples that fell outside the detection limit at varying dilutions. The progesterone EIA’s inter-assay coefficient of variation (COV) and intra-assay COV ranged from 2.7 -8.3% and 4.9 -7.6%, respectively. The mean inter-assay COV was 14.7% for the EI, and the mean intra-assay COV was 5.2% for the ELISA. We report progesterone values as nanograms per gram of blubber (ng g^1^). We repeated the extraction and measurements for a subset of the blubber samples, in which case we report the averaged resulting progesterone level, and ran multiple samples at several dilutions.

We assigned pregnancy status using previously described methods (Kershaw et al., 2020; Pallin et al., 2018). Briefly, we utilized previously established models developed from female humpback whales of known pregnancy status from the Gulf of Maine and the Gulf of St Lawrence, to assign the pregnancy status of sampled females based on their blubber progesterone concentrations. Previous studies successfully applied this modeling approach to other populations (e.g., Western Antarctic Peninsula (Pallin et al., 2018), Oceania (Riekkola et al., 2018)). We determined pregnancy rates as the number of pregnant females divided by the total number of assayed females for years in which at least five samples were available, i.e., in which sample size allowed for reasonably robust estimation.

### Statistical analysis

We checked whether the sex ratio deviated significantly from parity (1:1) for each region (northern Norway in winter, Barents Sea) using a two-tailed exact binomial test for the Barents Sea, and one-tailed test for Norway. We then tested whether the pregnancy rate differed between the summer (June and September) and winter season, using a Chi-squared test of independence. We used Quasi-binomial Generalised Linear Models (GLMs) to investigate variation in annual pregnancy rates between 2011 and 2018, and over the feeding season between June and February, using a “logit” link function to take into account overdispersion in the pregnancy rate data. Given the limited and variable biopsy sample sizes and the variability in pregnancy rate estimates, it was important to consider these data in the context of their power to detect significant changes over time. The power of the GLMs was estimated using the pwr.f2.test function in the pwr package (R version 3.6.2 (R Core Team, 2019)). The power to detect a trend in the pregnancy rate over the 8-year study period was 17.4%, and the power to detect a trend through the feeding season was 6.08%. Thus, the variability in pregnancy rate estimates makes the detection of significant temporal trends unlikely. We used a significance threshold of p <0.05 to determine significance in all statistical tests. Results are presented as mean ± standard deviation, unless otherwise noted.

## Results

### Photographic collections

In northern Norway, the total number of photo-identified humpback whales per winter ranged from a minimum of six individuals in the first year off Andøya (2010) to a maximum of 408 individuals in the 2015/16 season off Kvaløya (Figure 3 and S1 Table in supporting information). The peak in sightings occurred between November and January. The cumulative curve of identifications began to plateau after the winter of 2015/16 but showed a slight increase in 2018/19 in Kvænangen (Figure 3). In the Barents Sea, we registered humpback whale sightings from May to September, although most were photographed in September 2018. In total, we found five between-season re-sightings in the Barents Sea.

### Connectivity between Barents Sea and Norway

We matched 39 individual humpback whales between the Barents Sea and northern Norway (Figure 2). One individual was photographed in two different summers in the Barents Sea and subsequently re-sighted off northern Norway during winter both these years. All re-sightings occurred during the autumn and winter months in northern Norway. Among these 40 matches, 17 matches of 16 individuals occurred within the same season (Figure 2), most were first sighted in Norway in the end of November (S2 Table in supporting information).

**Figure 2.**
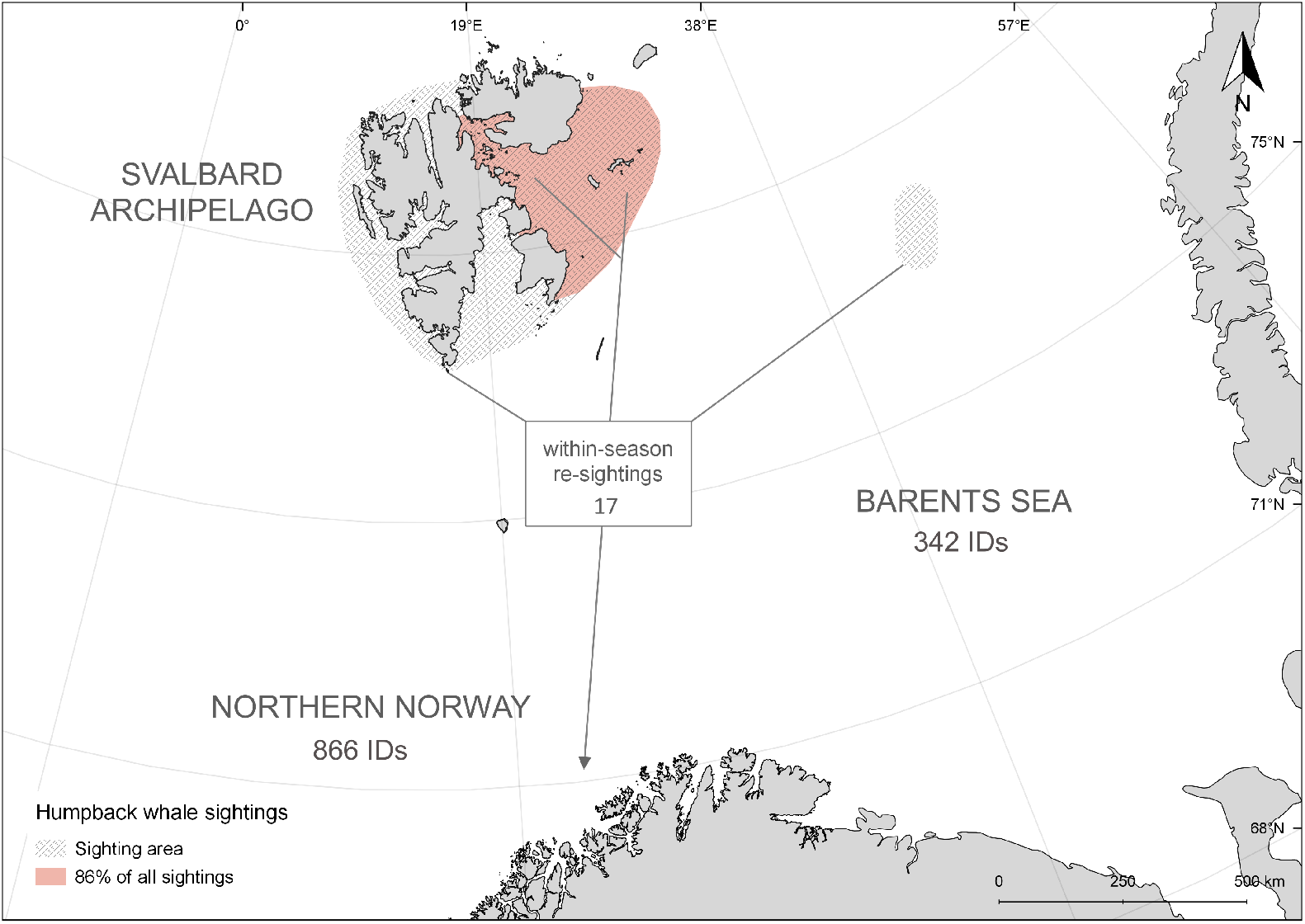
Map of sampling area in the Barents Sea. Map of sampling area in the Barents Sea (in grey and orange) with the number of identified individuals in the Barents Sea and northern Norway and the number of within-season matches. End of grey lines indicate locations from which individuals re-sighted in northern Norway were first reported. 86% of all humpback whale IDs in the Barents Sea were collected in the orange-shaded area.

**Figure 3.**
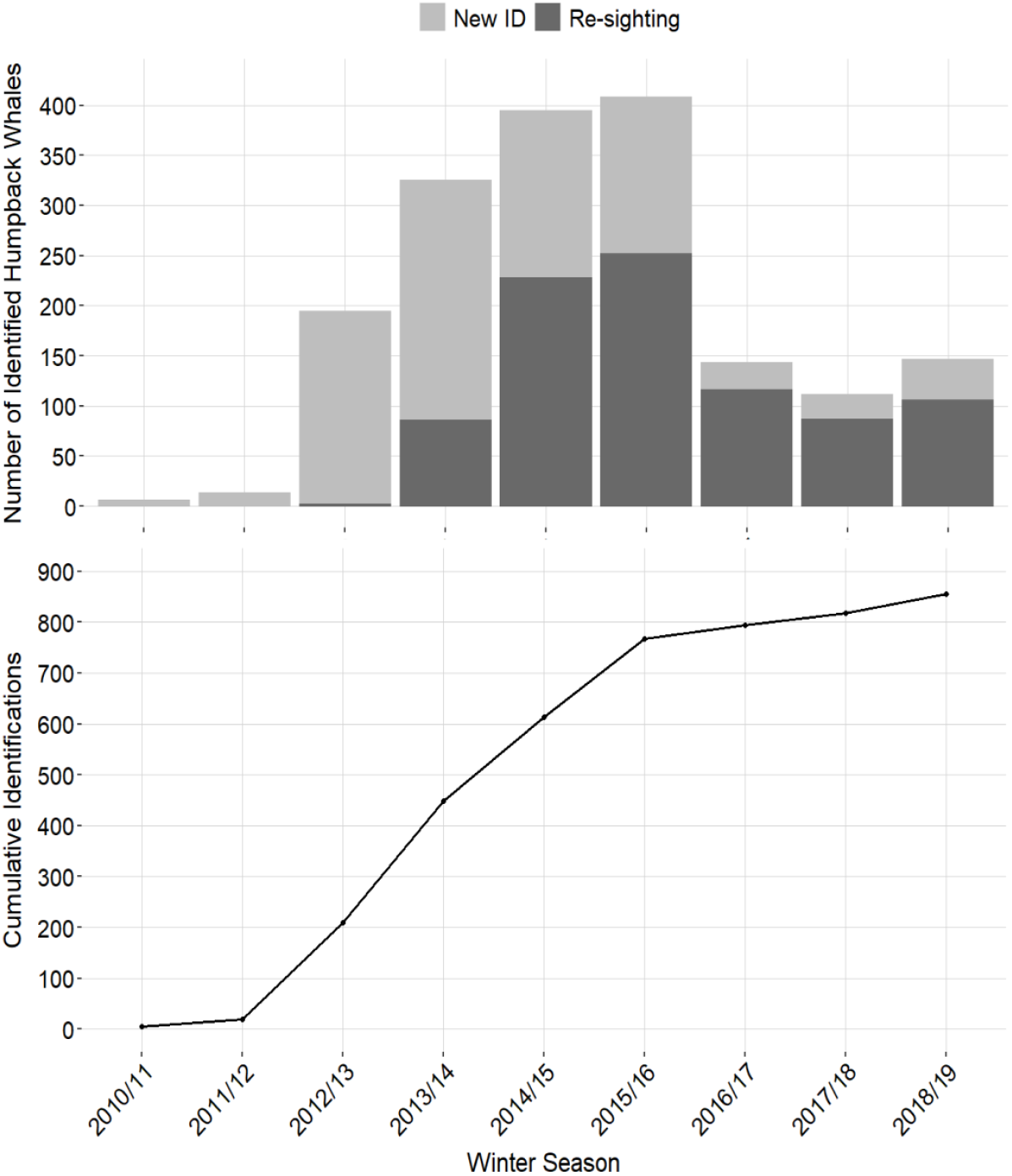
Re-sightings in Norway Upper panel: Total number of humpback whales (*Megaptera novaeangliae*) photo-identified each winter season in northern Norway, colored by new and re-sighted individuals from 2010/11 to 2018/19. Lower panel: Cumulative curve of photo-identified humpback whales during winter in northern Norway (2010/11 -2018/19).

### Site fidelity in northern Norway

Between the winter of 2010/11 and 2018/19, we photo-identified 866 individual humpback whales in northern Norway (Figure 3). The majority (53.4%, N = 457) returned in two or more winters. Most of these whales were seen in two (N = 202), three (N = 131) or four (N = 83) different years. The longest period over which an individual was re-sighted was seven years. Re-sightings between seasons occurred most frequently in sequential years (69.4%), followed by two-year intervals (20.6%) (Figure 4). Until the winter of 2013/14, new fluke captures accounted for more than 70% of the total number of whales identified in a season. In all following winters, the number of re-sightings was higher than first captures, on average 70.9% (± 10.5) (Figure 3).

**Figure 4.**
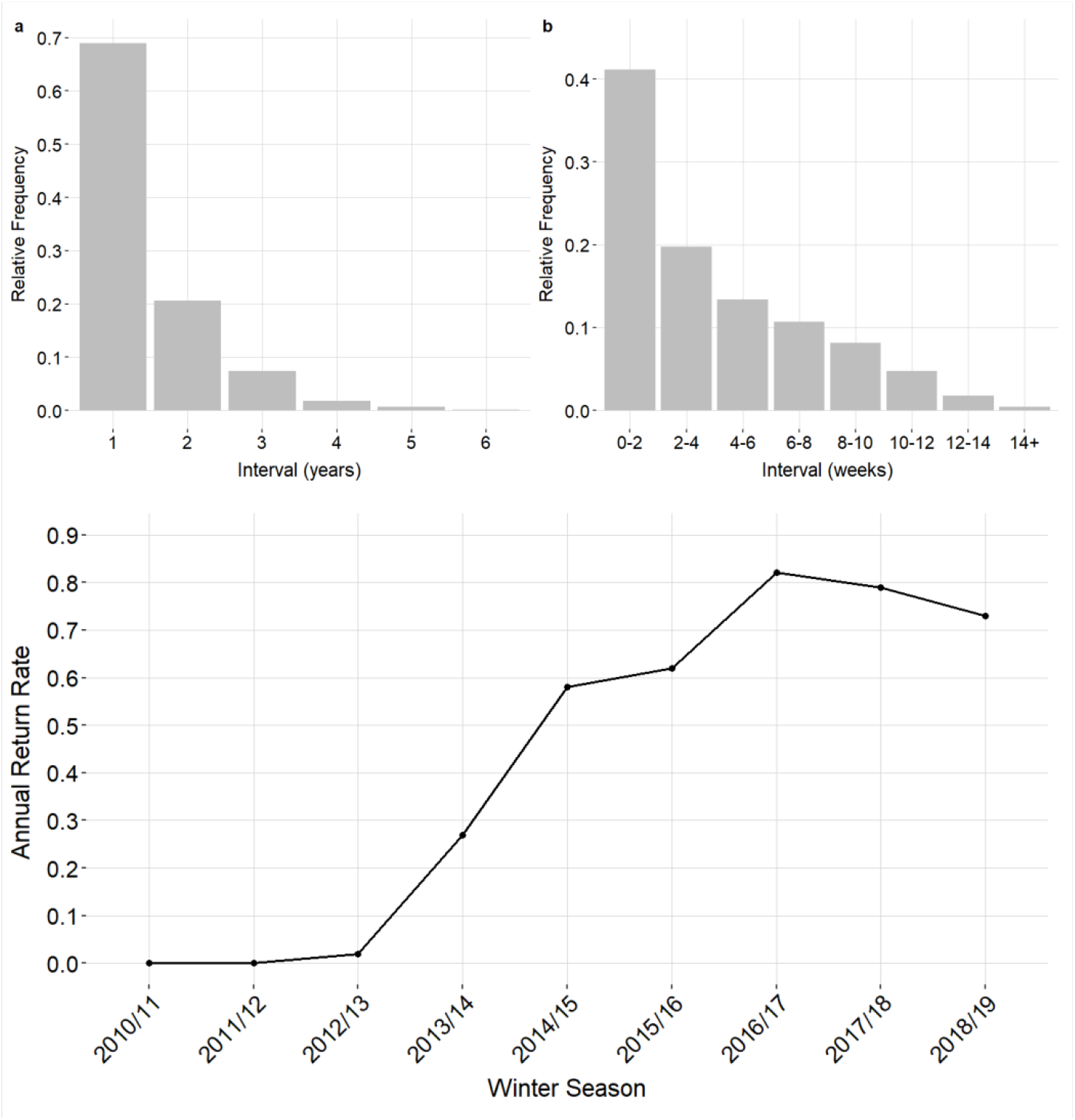
Numbers of identified humpback whales Upper panel: Between season re-sighting intervals (a) and within-season intervals (year seven not shown as it only contained one individual) (b) between first and last sighting of humpback whales (*Megaptera novaeangliae*) foraging off northern Norway during winter (2012/13 to 2018/19). Lower panel: Annual return rate of humpback whales to northern Norway (2010/11 to 2018/19).

The annual return rate, a measure of population-level site-fidelity, progressively increased until a peak in the 2016/17 season (the final winter season off Kvaløya, 81.8%; Figure 4), decreasing to 70 and 79% during the following two winter seasons (2017/18, 2018/19) in Kvænangen. 43.2% of the whales were seen more than once within a season. The time interval between within-season re-sightings ranged from a minimum of 2 days to a maximum of 15 weeks, on average 27.5 days (± 11.5; Figure 3). More than half the whales identified across the nine years of study were re-sighted, with 27% returning to feed for more than three years, most often in sequential years. In the winter of 2016/17, considerably fewer humpback whales were encountered around Kvaløya, and the first individuals were sighted in Kvænangen fjord. In the consecutive winter, the fjords around Kvaløya were deserted, and the feeding activity had shifted to Kvænangen fjord.

### Relatedness

A relatedness analysis based on a subset of the samples for which genetic sequences were available (107 individuals) indicated that no individuals were sampled repeatedly within the same season (coefficients of relatedness <1; S4 Table in supporting information).

### Sex ratio

The sex ratio in the Barents Sea was 1.4 (18M:13F, N= 31) and in northern Norway 0.6 (48M:76F, N= 124). No significant deviation from parity was found for the Barents Sea sample (p = 0.473), but the sex ratio differed significantly from parity in northern Norway with a bias in favor of females (p = 0.007). The sex ratio in northern Norway differed significantly between years in our sample (X^2^ = 12.9, p = 0.019), in years with low sample sizes (2011/12, 2017/18) the ratio of males in the sample was higher. The sex ratio did not differ significantly between months throughout the winter season (X^2^ = 3.2, p = 0.571; S1 Figure in supporting information).

### Pregnancy rate

All but three individuals were successfully assigned a reproductive status (i.e., pregnant or non-pregnant) by the reference model (with 99.9% confidence), all male controls were correctly classified as non-pregnant.

All progesterone concentrations are reported in S3 Table in supporting information. The pregnancy rate was low in the summer (22% northern Norway, June), and fall (20% Barents Sea, September 2018) and higher (median = 38%, 25^th^ quantile = 24%, 75^th^ quantile = 49%) during winter in northern Norway when pooled over all years (Table 2). However, the difference between the Barents Sea and Norway in 2018/19 (20% vs. 47%) was not statistically significant (X^2^ = 2, p = 1). Rates in winter varied across years between 8 -56% (Table 2). During the winter season, the pregnancy rate declined after a peak in December (73%), to 26% in January and 17% in February (Table 3). Due to the limited sample size and high variance, the power to detect a relationship in the pregnancy rate over winters in the eight-year study period was low (17.4%), and over the months during the feeding season even lower (6.1%). Thus, the variability in pregnancy rate estimates makes the detection of significant temporal trends unlikely.

**Table 2.** Numbers of female humpback whales assessed for progesterone levels and pregnancy rates in the Barents Sea and northern Norway by area and season. The pregnancy rate (pregnant females/all assayed females) is reported for months with at least five samples.

| Area | Season | Females | Pregnant | Not pregnant | Pregnancy rate (%) |
| --- | --- | --- | --- | --- | --- |
| northern Norway | June 2011 | 2 | 1 | 1 | - |
|  | June 2012 | 7 | 1 | 6 | 14 |
| Barents Sea | September 2018 | 10 | 2 | 8 | 20 |
| northern Norway | Winter 2013 / 2014 | 7 | 2 | 5 | 29 |
|  | Winter 2015 / 2016 | 12 | 1 | 11 | 8 |
|  | Winter 2016 / 2017 | 9 | 5 | 4 | 56 |
|  | Winter 2017 / 2018 | 2 | 0 | 2 | - |
|  | Winter 2018 / 2019 | 30 | 14 | 16 | 47 |
| **Total** |  | **79** | **26** | **53** |  |

**Table 3.** Numbers of female humpback whales assessed for progesterone levels and pregnancy rates in the Barents Sea and northern Norway by area and month. The pregnancy rate (pregnant females/all assayed females) is reported for months with at least five samples.

| Area | Month | Females | Pregnant | Not pregnant | Pregnancy rate (%) |
| --- | --- | --- | --- | --- | --- |
| Northern Norway | June | 9 | 2 | 7 | 22 |
| Barents Sea | September | 10 | 2 | 8 | 20 |
| Northern Norway | October | 2 | 0 | 2 | - |
|  | November | 14 | 6 | 8 | 43 |
|  | December | 11 | 8 | 3 | 73 |
|  | January | 27 | 7 | 20 | 26 |
|  | February | 6 | 1 | 5 | 17 |
| **Total** |  | **79** | **26** | **53** |  |

## Discussion

Within-season matches between the Barents Sea and northern Norway confirm that some Northeast Atlantic humpback whales continued their foraging season in northern Norwegian fjords. Females generally leave feeding grounds later than males resulting in a female bias late in the foraging season (Barendse et al., 2013; Dawbin, 1966; Pallin et al., 2018), consistent with our observation of a female bias in Norway but not the Barents Sea. Per our expectations, the pregnancy rate estimated during winter in northern Norway was higher than in June and September, indicating that these animals may indeed maximized their energy intake by continuing their foraging season in Norway. This was also observed in other areas (Pallin et al., 2018); summer 59%, fall 72%). The establishment of the foraging site in northern Norway coincided with dense herring concentrations in the area since 2010 (documented in detail since 2015 by e.g., (ICES, 2019)). The northward shift in the whales’ foraging aggregation over the years followed the shifting distribution of NSS-herring, which is most likely related to changes in the stock’s age structure (Huse et al., 2010). The high annual return rate, comparable to main feeding grounds (Calambokidis et al., 1996; Clapham et al., 1993; Ramp et al., 2010), indicates that this area has become an important part of the annual migration for some Northeast Atlantic humpback whales. Since the feeding activity is coupled to herring overwintering distribution, future shifts in the whales’ winter distribution can be expected as the herring changes its migration patterns and overwintering areas.

Information on the migration phenology of Northeast Atlantic humpback whales remains sparse due to the logistic challenges involved in surveying the Barents Sea region. Therefore, the duration of the foraging season is unknown. Sightings collected in the Barents Sea confirm that the area east of the Svalbard archipelago is an important foraging ground for humpback whales in late summer/autumn. This supports previous evidence from annual ecosystem surveys, whaling records, and tracking data (Hamilton et al., 2021; Nøttestad et al., 2015; van der Meeren & Prozorkevich, 2018). Unpublished tracking data from 2018 indicates that whales initiated migration from the Barents Sea between October and December in that year (Kettemer et al., 2022). Within-season resighting patterns in northern Norway show that most whales stayed longer than two weeks, many for about one month and some up to three months. Photographic matching to the breeding grounds in the West Indies (Stevick et al., 2018), along with a recently recorded round-trip migration by a female humpback whale (Kettemer et al., 2022) and unpublished tracking data show that animals can still migrate to breeding areas after foraging in northern Norway during the winter. However, pregnant females delaying their migration until late in the season may give birth along the migration route despite increasing their migration speed (Kettemer et al., 2022).

While our results indicate that pregnant females preferentially visit northern Norway as a continuation of the feeding season in the Barents Sea, we could not confirm that this group remained the longest in Norway. However, the statistical power to detect temporal trends in our data was low. One explanation for the lower pregnancy rates at the end of the season (January/February) may be that not all humpback whales complete migrations every year as shown in other feeding areas (Magnúsdóttir et al., 2019; Swingle et al., 1993). Juvenile individuals and resting females for whom the cost and risks outweigh the benefits of migration may therefore dominate the sample towards the end of the season, leading to low pregnancy rates among females sampled, as well as an increase in the proportion of males in February.

Monitoring pregnancy rates over time can indicate population health and growth rates, provided that the sample sizes are sufficient (Kershaw et al., 2020; Pallin et al., 2018). Our estimate of the variation in pregnancy rate between years is likely not sufficiently robust to infer trends in reproductive rates, due to the low number of samples in some years. Overall, the pregnancy rate in summer/fall and winter was lower than those reported on other foraging grounds. Elsewhere, pregnancy rates were reported to be higher, for example, 57% in the Southern Ocean (Riekkola et al., 2018), 58% (36-86%) in the Western Antarctic Peninsula (Pallin et al., 2018), 19 -48% in the North Pacific (Clark et al., 2016), and 25% -63% in the northwest Atlantic (Kershaw et al., 2020). The variability between years reported here was similarly high in those other studies. Pregnancy rate estimates present a minimum of true rates, as they usually include immature females. Pregnancy rates sampled at different times of the gestation period may vary, e.g., be inflated by subsequently aborted/reabsorbed pregnancies when sampled early, or missed if progesterone levels are not yet high enough (Pallin et al., 2018; Riekkola et al., 2018). However, recent work shows that pregnancy rates are tightly linked to fluctuations in prey availability (Pallin et al. in prep). Further studies should assess whether low rates here may indicate slowing population growth following recovery from exploitation (Leonard & Øien, 2020) or poor nutritional status due to changing environmental conditions (Wasser et al., 2017).

Rapid and fundamental ecosystem changes in the Barents Sea associated with warming, ice loss, and increased inflow of Atlantic waters have impacted a core foraging habitat of humpback whales (Hamilton et al., 2021; Johannesen et al., 2012). Generalist predators such as humpback whales are considered to be likely to adapt to changes in prey distributions (Nøttestad et al., 2015) and their populations have been increasing in recent decades after historical exploitation (Pallin et al., 2018; Zerbini et al., 2019). However, recently, humpback whale populations in the Northwest Atlantic and North Pacific have been experiencing declining calving rates, likely due to ecosystem shifts mediated by climate change (Cartwright et al., 2019; Kershaw et al., 2020). Climate change is projected to severely impact population vital rates and distributions of top predators on longer time scales (Descamps et al., 2017; Meyer-Gutbrod et al., 2015; Moore et al., 2019; Seyboth et al., 2016). Continued monitoring of the pregnancy or calving rate in this population is warranted as the ecosystems of the Barents and Norwegian seas shifts to a new ecological state. As anthropogenic use of the Arctic Ocean is increasing, knowledge of year-round distributions and critical habitat, especially during potentially vulnerable periods such as pregnancy, are essential for mitigating adverse effects of human activities on top predators.

## Conclusion

We confirm that humpback whales continued foraging in fjords of northern Norway during winter after their foraging season in the Barents Sea, and our results indicate that the area may be preferentially visited by pregnant females. Our findings suggest that winter foraging in northern Norway has become an important annual event for individual humpback whales, contingent on herring overwintering in these fjords. Over the last decades, the population of humpback whales in the Northeast Atlantic has recovered from historical over exploitation, while the Arctic ecosystem in which they forage is undergoing rapid and fundamental changes. The establishment of this foraging site is evidence of humpback whales’ ability to respond flexibly to prey resources along their migratory pathways, potentially affecting their migration phenology. Future work should aim to understand how this additional foraging opportunity impacts the overall fitness of individual whales and population vital rates.

## supporting information

**S1 Figure**

**Sex ratio by month** Sex ratio among humpback whales (*Megaptera novaeangliae*) sampled in the Barents Sea and northern Norway.

**S1 Table**

**Overview of photographic identification for each winter season in coastal Norway** Individuals identified, cumulative identifications, new individuals, re-sighted individuals, annual return rate and within-season re-sightings.

**S2 Table**

**Re-sighted individuals between the Barents Sea and northern Norway**. Sightings-ID, sighting locations and dates for individuals sighted both in the Barents Sea and Norway.

**S3 Table**

**Table of all progesterone values and model results**. Model results and progesterone levels for all samples run, including results of the subset of samples extracted with both methods for comparison.

**S4 Table**

**Pairwise coefficients of relatedness matrix** Resulting pairwise relatedness coefficients for a subset of individuals.

**S1 Text**

**Comparison of extraction methods** Details on the difference between the two different extractions used for progesterone analysis.

**S2 Text**

**Relatedness methods** Detailed methods for within-season re-sampling assessment using rxy.

## Acknowledgement

We are grateful to everyone who submitted pictures to NNHWC. We also thank Lars Kleivane and Kjell-Arne Fagerheim for their support during fieldwork. We thank Kim Praebel, Julie Bitz-Thorsen, and Shripathi Bhat for supporting the genetic analysis and providing laboratory space and materials. The project was supported by the “Whalefeast project” (financed by The Regional Research Council in Troms), and the UiT -The Arctic University of Norway (in Tromsø) and the Institute for Marine Research in Bergen.

## References

Abrahms, B., Hazen, E. L., Aikens, E. O., Savoca, M. S., Goldbogen, J. A., Bograd, S. J., Jacox, M. G., Irvine, L. M., Palacios, D. M., & Mate, B. R. (2019). Memory and resource tracking drive blue whale migrations. Proceedings of the National Academy of Sciences of the United States of America, 116(12), 5582–5587. https://doi.org/10.1073/pnas.1819031116

Avila, I. C., Dormann, C. F., García, C., Payán, L. F., & Zorrilla, M. X. (2020). Humpback whales extend their stay in a breeding ground in the Tropical Eastern Pacific. ICES Journal of Marine Science, 77 (1), 109–118. https://doi.org/10.1093/icesjms/fsz251

Barendse, J., Best, P. B., Carvalho, I., & Pomilla, C. (2013). Mother knows best: Occurrence and associations of resighted humpback whales suggest maternally derived fidelity to a Southern Hemisphere coastal feeding ground. PLoS ONE, 8(12). https://doi.org/10.1371/journal.pone.0081238

Bérubé, M., & Palsbøll, P. (1996). Identification of sex in Cetaceans by multiplexing with three ZFX and ZFY specific primers. Molecular Ecology, 5(2), 283–287. https://doi.org/10.1111/j.1365-294x.1996.tb00315.x

Calambokidis, J., Steiger, G. H., EvENsoN, J. R., Flynn, K. R., Balcomb, K. C., Claridge, D. E., Bloedel, P., Straley, J. M., Baker, C. S., & Ziegesar, O. V. (1996). Interchange and isolation of humpback whales off California and other North Pacific feeding grounds. Marine Mammal Science, 12(2), 215–226. https://doi.org/10.1111/j.1748-7692.1996.tb00572.x

Calderan, S. V., Black, A., Branch, T. A., Collins, M. A., Kelly, N., Leaper, R., Lurcock, S., Miller, B. S., Moore, M., Olson, P. A., irović, A., Wood, A. G., & Jackson, J. A. (2020). South Georgia blue whales five decades after the end of whaling. Endangered Species Research, 43, 359–373. https://doi.org/10.3354/esr01077

Cartwright, R., Venema, A., Hernandez, V., Wyels, C., Cesere, J., & Cesere, D. (2019). Fluctuating reproductive rates in Hawaii’s humpback whales, Megaptera novaeangliae, reflect recent climate anomalies in the North Pacific. Royal Society Open Science, 6(3). https://doi.org/10.1098/rsos.181463

Christensen, I., Haug, T., & Øien, N. (1992). Seasonal distribution, exploitation and present abundance of stocks of large baleen whales (Mysticeti) and sperm whales (Physeter macrocephalus) in Norwegian and adjacent waters [ISBN: 1054-3139]. ICES Journal of Marine Science, 49(3), 341–355. https://doi.org/10.1093/icesjms/49.3.341

Clapham, P. J., Baraff, L. S., Carlson, C. A., Christian, M. A., Mattila, D. K., Mayo, C. A., Murphy, M. A., & Pittman, S. (1993). Seasonal occurrence and annual return of humpback whales, Megaptera novaeangliae, in the southern Gulf of Maine. Can. J. Zool., 71, 440–443. https://doi.org/10.1139/z93-063

Clark, C. T., Fleming, A. H., Calambokidis, J., Kellar, N. M., Allen, C. D., Catelani, K. N., Robbins, M., Beaulieu, N. E., Steel, D., & Harvey, J. T. (2016). Heavy with child? Pregnancy status and stable isotope ratios as determined from biopsies of humpback whales [ISBN: 1474-6514]. Conservation Physiology, 4(1), 1–13. https://doi.org/10.1093/conphys/cow050

Davidson, S. C., Bohrer, G., Gurarie, E., LaPoint, S., Mahoney, P. J., Boelman, N. T., Eitel, J. U. H., Prugh, L. R., Vierling, L. A., Jennewein, J., Grier, E., Couriot, O., Kelly, A. P., Meddens, A. J. H., Oliver, R. Y., Kays, R., Wikelski, M., Aarvak, T., Ackerman, J. T., … Hebblewhite, M. (2020). Ecological insights from three decades of animal movement tracking across a changing Arctic. Science, 370(6517), 712–715. https://doi.org/10.1126/science.abb7080

Dawbin, W. H. (1966). The seasonal migratory cycle of humpback whales (K. S. Norris, Ed.) [Publication Title: Whales, Dolphins and Porpoises]. University of California Press. https://doi.org/10.1525/9780520321373-011

Descamps, S., Aars, J., Fuglei, E., Kovacs, K. M., Lydersen, C., Pavlova, O., Pedersen, Å. Ø., Ravolainen, V., & Strøm, H. (2017). Climate change impacts on wildlife in a High Arctic archipelago – Svalbard, Norway. Global Change Biology, 23(2), 490–502. https://doi.org/10.1111/gcb.13381

Hamilton, C., Lydersen, C., Aars, J., Biuw, M., Boltunov, A., Born, E., Dietz, R., Folkow, L., Glazov, D., Haug, T., Heide-Jørgensen, M., Kettemer, L., Laidre, K., Øien, N., Nordøy, E., Rikardsen, A., Rosing-Asvid, A., Semenova, V., Shpak, O., … Kovacs, K. (2021). Marine mammal hotspots in the Greenland and Barents Seas. Marine Ecology Progress Series, 659, 3–28. https://doi.org/10.3354/meps13584

Hanghøj, K., Moltke, I., Andersen, P. A., Manica, A., & Korneliussen, T. S. (2019). Fast and accurate relatedness estimation from high-throughput sequencing data in the presence of inbreeding. GigaScience, 8(5), giz034. https://doi.org/10.1093/gigascience/giz034

Huse, G., Fernö, A., & Holst, J. C. (2010). Establishment of new wintering areas in herring co-occurs with peaks in the ‘first time/repeat spawner’ ratio. Marine Ecology Progress Series, 409, 189–198. https://doi.org/10.3354/meps08620

ICES. (2019). Working Group on Widely Distributed Stocks (WGWIDE) (Rep.). ICES Scientific Reports. https://doi.org/10.17895/ices.pub.5574

Ingebrigtsen, A. (1929). Whales caught in the North Atlantic and other seas (Rapport et procès-verbaux des réunions No. 56). Conseil Permanent International pour l’Exploration de la Mer.

Johannesen, E., Ingvaldsen, R. B., Bogstad, B., Dalpadado, P., Eriksen, E., Gjøsæter, H., Knutsen, T., Skern-Mauritzen, M., & Stiansen, J. E. (2012). Changes in Barents Sea ecosystem state, 19702009: Climate fluctuations, human impact, and trophic interactions. ICES Journal of Marine Science, 69(5), 880–889. https://doi.org/10.1093/icesjms/fss046

Jourdain, E., & Vongraven, D. (2017). Humpback whale (Megaptera novaeangliae) and killer whale (Orcinus orca) feeding aggregations for foraging on herring (Clupea harengus) in Northern Norway. Mammalian Biology, 86, 27–32. https://doi.org/10.1016/j.mambio.2017.03.006

Katona, S. K., & Whitehead, H. P. (1981). Identifying humpback whales using their natural markins. Polar Record, 20(128), 439–444. https://doi.org/10.1017/s003224740000365x

Keen, E. M., Pilkington, J., O’Mahony, É., Thompson, K.-L., Hendricks, B., Robinson, N., Dundas, A., Nichol, L., Alidina, H. M., Meuter, H., Picard, C. R., & Wray, J. (2021). Fin whales of the Great Bear Rainforest: Balaenoptera physalus velifera in a Canadian Pacific fjord system. PLOS ONE, 16(9), e0256815. https://doi.org/10.1371/journal.pone.0256815

Kershaw, J. L., Ramp, C. A., Sears, R., Plourde, S., Brosset, P., Miller, P. J. O., & Hall, A. J. (2020). Declining reproductive success in the Gulf of St. Lawrence’s humpback whales (Megaptera novaeangliae) reflects ecosystem shifts on their feeding grounds. Global Change Biology, 00(00), 1–15. https://doi.org/10.1111/gcb.15466

Kettemer, L. E., Rikardsen, A. H., Biuw, M., Broms, F., Mul, E., & Blanchet, M.-A. (2022). Round-trip migration and energy budget of a breeding female humpback whale in the Northeast Atlantic. PLOS ONE, 17 (5), e0268355. https://doi.org/10.1371/journal.pone.0268355

Korneliussen, T. S., Albrechtsen, A., & Nielsen, R. (2014). ANGSD: Analysis of Next Generation Sequencing Data. BMC Bioinformatics, 15(1), 1–13. https://doi.org/10.1186/s12859-014-0356-4

Kramvig, B., Kristoffersen, B., & Førde, A. (2016). Responsible Cohabitation in Arctic Waters: The Promise of a Spectacle Tourist Whale. In S. Abram & K. A. Lund (Eds.), Green Ice: Tourism Ecologies in the European High North (pp. 25–47). Palgrave Macmillan UK. https://doi.org/10.1057/978-1-137-58736-7_2

Leonard, D., & Øien, N. (2020). Estimated abundances of cetacean species in the Northeast Atlantic from Norwegian shipboard surveys conducted in 2014–2018. NAMMCO Scientific Publications, 11. https://doi.org/10.7557/3.4694

Magnúsdóttir, E. E., Víkingsson, G. A., Hali, A., Whittaker, M., Chosson, V., Pampoulie, C., Svavarsson, J., & Miller, P. (2019). Subarctic winter whales: An overwintering strategy of humpback whales in Icelandic waters [Presented at: World Marine Mammal Conference]. https://doi.org/10/gqx295

Meredith, M., Sommerkorn, M., Cassotta, S., Derksen, C., Ekaykin, A., Hollowed, A., Kofinas, G., Mackintosh, A., Melbourne-Thomas, J., Muelbert, M., Ottersen, G., Pritchard, H., & Schuur, E. (2019). Polar Regions. In H.-O. Pörtner, D.C. Roberts, V. Masson-Delmotte, P. Zhai, M. Tignor, E. Poloczanska, K. Mintenbeck, A. Alegría, M. Nicolai, A. Okem, J. Petzold, B. Rama, N.M. Weyer (Ed.), IPCC Special Report on the Ocean and Cryosphere in a Changing Climate (pp. 203–320). Cambridge University Press. https://doi.org/10.1017/9781009157964.005

Meyer-Gutbrod, E. L., Greene, C. H., Sullivan, P. J., & Pershing, A. J. (2015). Climate-associated changes in prey availability drive reproductive dynamics of the North Atlantic right whale population. Marine Ecology Progress Series, 535, 243–258. https://doi.org/10.3354/meps11372

Moore, S. E., Haug, T., Víkingsson, G. A., & Stenson, G. B. (2019). Baleen whale ecology in arctic and subarctic seas in an era of rapid habitat alteration. Progress in Oceanography, 176, 102118. https://doi.org/10.1016/j.pocean.2019.05.010

Nøttestad, L., Krafft, B. A., Anthonypillai, V., Bernasconi, M., Langård, L., Mørk, H. L., & Fernö¶, A. (2015). Recent changes in distribution and relative abundance of cetaceans in the Norwegian Sea and their relationship with potential prey. Frontiers in Ecology and Evolution, 2, 83. https://doi.org/10.3389/fevo.2014.00083

Pallin, L. J., Baker, C. S., Steel, D., Kellar, N. M., Robbins, J., Johnston, D. W., Nowacek, D. P., Read, A. J., & Friedlaender, A. S. (2018). High pregnancy rates in humpback whales (Megaptera novaeangliae) around the Western Antarctic Peninsula, evidence of a rapidly growing population. Royal Society Open Science, 5(5). https://doi.org/10.1098/rsos.180017

R Core Team. (2019). A language and environment for statistical computing. [URL: http://www.R-project.org/.].

Ramp, C., Bérubé, M., Palsbøll, P., Hagen, W., & Sears, R. (2010). Sex-specific survival in the humpback whale Megaptera novaeangliae in the Gulf of St. Lawrence, Canada. Marine Ecology Progress Series, 400, 267–276. https://doi.org/10.3354/meps08426

Ramp, C., Delarue, J., Palsbøll, P. J., Sears, R., & Hammond, P. S. (2015). Adapting to a warmer ocean - Seasonal shift of baleen whale movements over three decades. PLoS ONE, 10(3), 1–15. https://doi.org/10.1371/journal.pone.0121374

Riekkola, L., Andrews-Goff, V., Friedlaender, A., Constantine, R., Zerbini, A. N., Andrews, O., Andrews-Goff, V., Baker, C. S., Chandler, D., Childerhouse, S., Clapham, P., Dodémont, R., Donnelly, D., Friedlaender, A., Gallego, R., Garrigue, C., Ivashchenko, Y., Jarman, S., Lindsay, R., … Constantine, R. (2018). Application of a multi-disciplinary approach to reveal population structure and Southern Ocean feeding grounds of humpback whales. Ecological Indicators, 89, 455–465. https://doi.org/10.1016/j.ecolind.2018.02.030

Seyboth, E., Groch, K. R., Dalla Rosa, L., Reid, K., Flores, P. A. C., & Secchi, E. R. (2016). Southern Right Whale (Eubalaena australis) Reproductive Success is Influenced by Krill (Euphausia superba) Density and Climate. Scientific Reports, 6(1), 28205. https://doi.org/10.1038/srep28205

Stevick, P. T., Bouveret, L., Gandilhon, N., Rinaldi, C., Broms, F., Carlson, C., Kennedy, A., Ward, N., & Wenzel, F. (2018). Migratory destinations and timing of humpback whales in the southeastern Caribbean differ from those off the Dominican Republic. J Cetacean Res. Manage., 18, 127–133.

Swingle, W. M., Barco, S. G., Pitchford, T. D., Mclellan, W. A., & Pabst, D. A. (1993). Appearance of Juvenile Humpback Whales Feeding in the Nearshore Waters of Virginia [ISBN: 0824-0469]. Marine Mammal Science, 9(3), 309–315. https://doi.org/10.1111/j.1748-7692.1993.tb00458.x

van der Meeren, G. i., & Prozorkevich, D. (2018). Survey report from the joint Norwegian/Russian Ecosystem Survey in the Barents Sea and adjacent waters, August–October 2017 (tech. rep. No. 2-2018). IMR-PINRO.

Wasser, S. K., Lundin, J. I., Ayres, K., Seely, E., Giles, D., Balcomb, K., Hempelmann, J., Parsons, K., & Booth, R. (2017). Population growth is limited by nutritional impacts on pregnancy success in endangered Southern Resident killer whales (Orcinus orca). PLOS ONE, 12(6), e0179824. https://doi.org/10.1371/journal.pone.0179824

Wenzel, F. W., Broms, F., López-Suárez, P., Lopes, K., Veiga, N., Yeoman, K., Rodrigues, M. S. D., Allen, J., Fernald, T. W., Stevick, P. T., Jones, L., Jann, B., Bouveret, L., Ryan, C., Berrow, S., & Corkeron, P. (2020). Humpback whales (Megaptera novaeangliae) in the Cape Verde Islands: Migratory patterns, resightings, and abundance. Aquatic Mammals, 46(1), 21–31. https://doi.org/10.1578/am.46.1.2020.21

Zerbini, A. N., Adams, G., Best, J., Clapham, P. J., Jackson, J. A., & Punt, A. E. (2019). Assessing the recovery of an Antarctic predator from historical exploitation. Royal Society Open Science, 6(10), 190368. https://doi.org/10.1098/rsos.190368

