## supporting information for "Arctic humpback whales respond to nutritional opportunities before migration"

**Figure S1 Sex ratio by month.** Sex ratio among humpback whales (*Megaptera novaeangliae*) sampled in the Barents Sea and northern Norway.

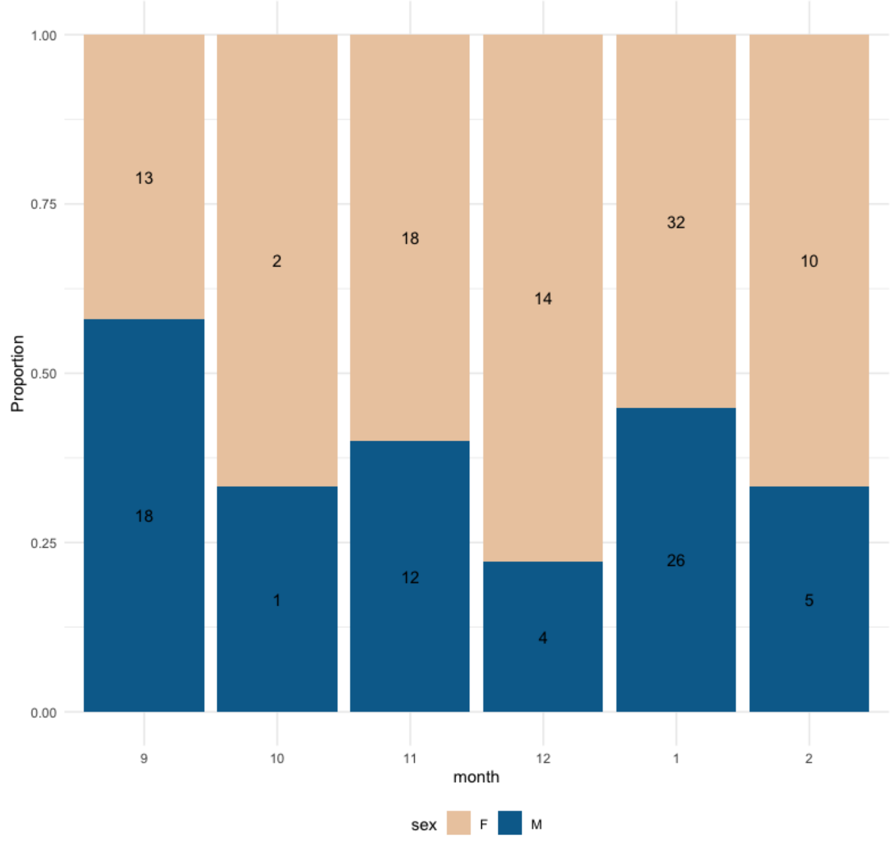

**Table S1 Overview of photographic identification for each winter season in coastal Norway.** Individuals identified, cumulative identifications, new individuals, re-sighted individuals, annual return rate and within-season re-sightings.

| Year | 2010/11 | 2011/12 | 2012/13 | 2013/14 | 2014/15 | 2015/16 | 2016/17 | 2017/18 | 2018/19 |
|---|---|---|---|---|---|---|---|---|---|
| Total no. of identified individuals/year | 6 | 13 | 194 | 325 | 394 | 408 | 143 | 111 | 146 |
| Cumulative identifications | 6 | 19 | 210 | 448 | 613 | 768 | 794 | 817 | 856 |
| No. of new individuals/year | 6 | 13 | 191 | 238 | 165 | 155 | 26 | 23 | 39 |
| No. of re-sighted individuals/year | 0 | 0 | 3 | 87 | 229 | 253 | 117 | 88 | 107 |
| Annual return (%) | 0 | 0 | 2 | 27 | 58 | 62 | 82 | 79 | 73 |
| No. of within-season re-sights | 0 | 0 | 51 | 117 | 192 | 180 | 71 | 45 | 83 |

**Table S2 Re-sighted individuals between the Barents Sea and northern Norway.** Sightings-ID, sighting locations and dates for individuals sighted both in the Barents Sea and Norway.

| ID | summer feeding location | summer sighting date (m/d/y) | winter feeding location | winter first sighting | winter last sighting |
|---|---|---|---|---|---|
| NNHWC-117 | NE-Svalbard | 7/19/12 | Kvaløya | 12/9/12 | 1/2/13 |
| NNHWC-138 | NE-Svalbard | 8/25/12 | Kvaøya | 12/24/12 | 12/4/12 |
| NNHWC-139 | NE-Svalbard | 8/25/12 | Kvaøya | 11/27/12 | |
| NNHWC-156 | NE-Svalbard | 9/8/18 | Skjervøy | 10/28/18 | 12/19/18 |
| NNHWC-193 | Hinlopen Strait | 7/12/13 | Kvaløya | 12/2/13 | |
| NNHWC-286 | NE-Svalbard | 9/8/18 | Skjervøy | 11/13/18 | 1/4/19 |
| NNHWC-286 | Border to Russia | 8/8/15 | Kvaløya | 11/17/15 | |
| NNHWC-295 | Hinlopen Strait | 7/7/14 | Kvaløya | 11/16/14 | 01.12.1014 |
| NNHWC-344 | NE-Svalbard | 9/5/18 | Skjervøy | 12/17/18 | |
| NNHWC-387 | Hinlopen Strait | 7/7/14 | Kvaløya | 12/21/14 | |
| NNHWC-471 | Hornsund | 8/28/18 | Skjervøy | 1/9/19 | |
| NNHWC-567 | Border to Russia | 8/8/15 | Kvaløya | 11/30/15 | 12/2/15 |
| NNHWC-609 | NE-Svalbard | 9/5/18 | Skjervøy | 12/18/18 | 1/9/19 |
| NNHWC-698 | NE-Svalbard | 9/3/18 | Skjervøy | 11/16/18 | 12/11/18 |
| T2-73 | NE-Svalbard | 9/5/18 | Skjervøy | 11/14/18 | 11/15/18 |
| T3-18 | NE-Svalbard | 9/4/18 | Skjervøy | 12/21/18 | |
| TT2-29 | NE-Svalbard | 9/9/18 | Skjervøy | 12/31/18 | |

**Table S3 All progesterone values and model results.** Model results and progesterone levels for all samples run, including results of the subset of samples extracted with both methods for comparison.

Table_S3

| Sample | Extraction | Year | Date | P4 | log(P4) | Preg | Pregnant | Month | Season | Notes | Probabil | Hi | Hi Correct | Low | Low Correct | Control (Male) | Re-extract |
|---|---|---|---|---|---|---|---|---|---|---|---|---|---|---|---|---|---|
| **Mnova16002** | 2016.01.024 | 2016 | 1/13/2016 | 0.956803476 | -0.019177256 | 0 | No | 1 | 15/16 | Coastal Norway | 2.22E-16 | 2.22E-16 | 0.00E+00 | 2.22E-16 | 0.00E+00 | | |
| **2016.01.019** | | 2016 | 2/6/2016 | 1.311666243 | 0.117823342 | 0 | No | 2 | 15/16 | Coastal Norway | 2.22E-16 | 2.22E-16 | 0.00E+00 | 2.22E-16 | 0.00E+00 | | |
| **Mnova16003** | 2016.01.008 | 2016 | 2/1/2016 | 0.557348336 | -0.253873291 | 0 | No | 2 | 15/16 | Coastal Norway | 2.22E-16 | 2.22E-16 | 0.00E+00 | 2.22E-16 | 0.00E+00 | | |
| **Mnova16006** | 2016.01.011 | 2016 | 2/1/2016 | 0.593032505 | -0.226921502 | 0 | No | 2 | 15/16 | Coastal Norway | 2.22E-16 | 2.22E-16 | 0.00E+00 | 2.22E-16 | 0.00E+00 | | |
| **Mnova16010** | 2016.01.015 | 2016 | 2/6/2016 | 198.8373512 | 2.298497969 | 1 | Yes | 2 | 15/16 | Coastal Norway | 1 | 1 | 0.00E+00 | 1 | 0.00E+00 | | |
| **Mnova16012** | 2016.01.017 | 2016 | 2/6/2016 | 0.622019928 | -0.206195702 | 0 | No | 2 | 15/16 | Coastal Norway | 2.22E-16 | 2.22E-16 | 0.00E+00 | 2.22E-16 | 0.00E+00 | | |
| **Mnova16014** | 2016.01.019 | 2016 | 2/6/2016 | 0.652813886 | -0.185210617 | 0 | No | 2 | 15/16 | Coastal Norway | 2.22E-16 | 2.22E-16 | 0.00E+00 | 2.22E-16 | 0.00E+00 | | |
| **Mnova16016** | 2016.01.021 | 2016 | 2/20/2016 | 0.69174374 | -0.160054762 | 0 | No | 2 | 15/16 | Coastal Norway | 2.22E-16 | 2.22E-16 | 0.00E+00 | 2.22E-16 | 0.00E+00 | | |
| **2016.01.006** | | 2016 | 11/7/2016 | 102.6236327 | 2.011247384 | 1 | Yes | 11 | 16/17 | Coastal Norway | 1 | 1 | 0.00E+00 | 1 | 0.00E+00 | | |
| **2016.01.001** | | 2016 | 12/31/2016 | 127.6640883 | 2.106068748 | 1 | Yes | 12 | 16/17 | Coastal Norway | 1 | 1 | 0.00E+00 | 1 | 0.00E+00 | | |
| **2017.01.007** | | 2017 | 1/4/2017 | 1.10359132 | 0.042808276 | 0 | No | 1 | 16/17 | Coastal Norway | 2.22E-16 | 2.22E-16 | 0.00E+00 | 2.22E-16 | 0.00E+00 | 1 | |
| **2017.01.008** | | 2017 | 1/4/2017 | 1.315398485 | 0.119057337 | 0 | No | 1 | 16/17 | Coastal Norway | 2.22E-16 | 2.22E-16 | 0.00E+00 | 2.22E-16 | 0.00E+00 | | |
| **2017.01.010** | | 2017 | 1/15/2017 | 164.7148019 | 2.216732628 | 1 | Yes | 1 | 16/17 | Coastal Norway | 1.00E+00 | 1.00E+00 | 0.00E+00 | 1.00E+00 | 0.00E+00 | | |
| **2017.01.011** | | 2017 | 1/13/2017 | 1.201915112 | 0.079873796 | 0 | No | 1 | 16/17 | Coastal Norway | 2.22E-16 | 2.22E-16 | 0.00E+00 | 2.22E-16 | 0.00E+00 | 1 | |
| **2017.01.012** | | 2017 | 1/13/2017 | 0.578473709 | -0.237716375 | 0 | No | 1 | 16/17 | Coastal Norway | 2.22E-16 | 2.22E-16 | 0.00E+00 | 2.22E-16 | 0.00E+00 | | |
| **2017.01.015** | | 2017 | 1/11/2017 | 6.773915265 | 0.83083976 | 0 | No | 1 | 16/17 | Coastal Norway | 2.47E-15 | 1.81E-06 | 1.81E-06 | 2.22E-16 | 2.25E-15 | | |
| **2017.01.001** | | 2017 | 11/14/2017 | 0.706203896 | -0.151069891 | 0 | No | 11 | 17/18 | Coastal Norway | 2.22E-16 | 2.22E-16 | 0.00E+00 | 2.22E-16 | 0.00E+00 | 1 | |
| **2017.01.005** | | 2017 | 11/14/2017 | 0.417354288 | -0.379495121 | 0 | No | 11 | 17/18 | Coastal Norway | 2.22E-16 | 2.22E-16 | 0.00E+00 | 2.22E-16 | 0.00E+00 | | |
| **2017.01.006** | | 2017 | 11/14/2017 | 0.843229911 | -0.074053997 | 0 | No | 11 | 17/18 | Coastal Norway | 2.22E-16 | 2.22E-16 | 0.00E+00 | 2.22E-16 | 0.00E+00 | 1 | |
| **2018.01.001** | | 2018 | 9/4/2018 | 0.718404075 | -0.143631213 | 0 | No | 9 | 18 | Barents Sea | 2.22E-16 | 2.22E-16 | 0.00E+00 | 2.22E-16 | 0.00E+00 | | |
| **2018.01.002** | | 2018 | 9/4/2018 | 1.109227287 | 0.045020544 | 0 | No | 9 | 18 | Barents Sea | 2.22E-16 | 2.22E-16 | 0.00E+00 | 2.22E-16 | 0.00E+00 | | |
| **2018.01.003** | | 2018 | 9/4/2018 | 303.7914581 | 2.482575558 | 1 | Yes | 9 | 18 | Barents Sea | 1 | 1 | 0.00E+00 | 1 | 0.00E+00 | | |
| **2018.01.004** | | 2018 | 9/4/2018 | 194.7449783 | 2.289466268 | 1 | Yes | 9 | 18 | Barents Sea | 1 | 1 | 0.00E+00 | 1 | 0.00E+00 | | |
| **2018.01.005** | | 2018 | 9/4/2018 | 0.882724529 | -0.054174805 | 0 | No | 9 | 18 | Barents Sea | 2.22E-16 | 2.22E-16 | 0.00E+00 | 2.22E-16 | 0.00E+00 | 1 | |
| **2018.01.009** | | 2018 | 9/7/2018 | 0.312042263 | -0.505786581 | 0 | No | 9 | 18 | Barents Sea | 2.22E-16 | 2.22E-16 | 0.00E+00 | 2.22E-16 | 0.00E+00 | | |
| **2018.01.015** | | 2018 | 9/7/2018 | 0.372677288 | -0.428667073 | 0 | No | 9 | 18 | Barents Sea | 2.22E-16 | 2.22E-16 | 0.00E+00 | 2.22E-16 | 0.00E+00 | 1 | |
| **2018.01.016** | | 2018 | 9/7/2018 | 0.486273738 | -0.313119185 | 0 | No | 9 | 18 | Barents Sea | 2.22E-16 | 2.22E-16 | 0.00E+00 | 2.22E-16 | 0.00E+00 | 1 | |
| **2018.01.017** | | 2018 | 9/8/2018 | 0.437398231 | -0.359122978 | 0 | No | 9 | 18 | Barents Sea | 2.22E-16 | 2.22E-16 | 0.00E+00 | 2.22E-16 | 0.00E+00 | | |
| **2018.01.018** | | 2018 | 9/8/2018 | 0.482736622 | -0.316289753 | 0 | No | 9 | 18 | Barents Sea | 2.22E-16 | 2.22E-16 | 0.00E+00 | 2.22E-16 | 0.00E+00 | | |
| **2018.01.020** | | 2018 | 9/8/2018 | 0.32306122 | -0.490715171 | 0 | No | 9 | 18 | Barents Sea | 2.22E-16 | 2.22E-16 | 0.00E+00 | 2.22E-16 | 0.00E+00 | 1 | |
| **2018.01.022** | | 2018 | 9/9/2018 | 0.324596969 | -0.48865554 | 0 | No | 9 | 18 | Barents Sea | 2.22E-16 | 2.22E-16 | 0.00E+00 | 2.22E-16 | 0.00E+00 | | |
| **2018.01.023** | | 2018 | 9/9/2018 | 0.168455488 | -0.773514836 | 0 | No | 9 | 18 | Barents Sea | 2.22E-16 | 2.22E-16 | 0.00E+00 | 2.22E-16 | 0.00E+00 | 1 | |
| **2018.01.026** | | 2018 | 9/9/2018 | 0.20848949 | -0.680915833 | 0 | No | 9 | 18 | Barents Sea | 2.22E-16 | 2.22E-16 | 0.00E+00 | 2.22E-16 | 0.00E+00 | | |
| **2018.01.028** | | 2018 | 9/9/2018 | 0.567278686 | -0.246203534 | 0 | No | 9 | 18 | Barents Sea | 2.22E-16 | 2.22E-16 | 0.00E+00 | 2.22E-16 | 0.00E+00 | | |
| **2018.01.032** | | 2018 | 10/26/2018 | 0.520011923 | -0.283986699 | 0 | No | 10 | 18/19 | Coastal Norway | 2.22E-16 | 2.22E-16 | 0.00E+00 | 2.22E-16 | 0.00E+00 | | |
| **2018.01.033** | | 2018 | 10/26/2018 | 0.609093895 | -0.215315753 | 0 | No | 10 | 18/19 | Coastal Norway | 2.22E-16 | 2.22E-16 | 0.00E+00 | 2.22E-16 | 0.00E+00 | | |
| **2018.01.038** | | 2018 | 11/6/2018 | 347.9784932 | 2.541552403 | 1 | Yes | 11 | 18/19 | Coastal Norway | 1 | 1 | 0.00E+00 | 1 | 0.00E+00 | | |
| **2018.01.039** | Mnova18059 | 2018 | 11/6/2018 | 0.91413743 | -0.038988508 | 0 | No | 11 | 18/19 | Coastal Norway | 2.22E-16 | 2.22E-16 | 0.00E+00 | 2.22E-16 | 0.00E+00 | | |
| **2018.01.040** | Mnova18060 | 2018 | 11/6/2018 | 0.456686488 | -0.340381838 | 0 | No | 11 | 18/19 | Coastal Norway | 2.22E-16 | 2.22E-16 | 0.00E+00 | 2.22E-16 | 0.00E+00 | | |
| **2018.01.041** | | 2018 | 11/6/2018 | 0.510225088 | -0.292238191 | 0 | No | 11 | 18/19 | Coastal Norway | 2.22E-16 | 2.22E-16 | 0.00E+00 | 2.22E-16 | 0.00E+00 | | |
| **2018.01.043** | Mnova18063 | 2018 | 11/7/2018 | 1.843337596 | 0.265604881 | 0 | No | 11 | 18/19 | Coastal Norway | 2.22E-16 | 2.22E-16 | 0.00E+00 | 2.22E-16 | 0.00E+00 | 1 | |
| **2018.01.044** | Mnova18064 | 2018 | 11/7/2018 | 79.58533376 | 1.900833042 | 1 | Yes | 11 | 18/19 | Coastal Norway | 1.00E+00 | 1.00E+00 | 0.00E+00 | 1.00E+00 | 0.00E+00 | | |
| **2018.01.045** | | 2018 | 11/7/2018 | 153.6396624 | 2.186503344 | 1 | Yes | 11 | 18/19 | Coastal Norway | 1.00E+00 | 1.00E+00 | 0.00E+00 | 1.00E+00 | 0.00E+00 | | |
| **2018.01.046** | | 2018 | 11/7/2018 | 1.933410686 | 0.286324115 | 0 | No | 11 | 18/19 | Coastal Norway | 2.22E-16 | 2.22E-16 | 0.00E+00 | 2.22E-16 | 0.00E+00 | 1 | |
| **2018.01.047** | | 2018 | 11/7/2018 | 261.1418481 | 2.416876473 | 1 | Yes | 11 | 18/19 | Coastal Norway | 1.00E+00 | 1.00E+00 | 0.00E+00 | 1.00E+00 | 0.00E+00 | | |
| **2018.01.048** | | 2018 | 11/13/2018 | 0.655320773 | -0.183546065 | 0 | No | 11 | 18/19 | Coastal Norway | 2.22E-16 | 2.22E-16 | 0.00E+00 | 2.22E-16 | 0.00E+00 | | |
| **2018.01.049** | | 2018 | 11/13/2018 | 0.383548177 | -0.416180077 | 0 | No | 11 | 18/19 | Coastal Norway | 2.22E-16 | 2.22E-16 | 0.00E+00 | 2.22E-16 | 0.00E+00 | | |
| **2018.01.051** | | 2018 | 11/15/2018 | 872.4925177 | 2.940761711 | 1 | Yes | 11 | 18/19 | Coastal Norway | 1 | 1 | 0.00E+00 | 1 | 0.00E+00 | | |
| **2018.01.053** | Mnova18072 | 2018 | 12/2/2018 | 190.3375529 | 2.279524482 | 1 | Yes | 12 | 18/19 | Coastal Norway | 1 | 1 | 0.00E+00 | 1 | 0.00E+00 | | |
| **2018.01.054** | Mnova18073 | 2018 | 12/2/2018 | 0.811878619 | -0.090508896 | 0 | No | 12 | 18/19 | Coastal Norway | 2.22E-16 | 2.22E-16 | 0.00E+00 | 2.22E-16 | 0.00E+00 | | |
| **2018.01.057** | | 2018 | 12/4/2018 | 136.0270604 | 2.133625313 | 1 | Yes | 12 | 18/19 | Coastal Norway | 1 | 1 | 0.00E+00 | 1 | 0.00E+00 | | |
| **2018.01.058** | | 2018 | 12/4/2018 | 106.6856462 | 2.028105992 | 1 | Yes | 12 | 18/19 | Coastal Norway | 1 | 1 | 0.00E+00 | 1 | 0.00E+00 | | |
| **2018.01.059** | | 2018 | 12/4/2018 | 0.925390389 | -0.033675016 | 0 | No | 12 | 18/19 | Coastal Norway | 2.22E-16 | 2.22E-16 | 0.00E+00 | 2.22E-16 | 0.00E+00 | | |
| **2018.01.060** | | 2018 | 12/4/2018 | 84.05189925 | 1.924547531 | 1 | Yes | 12 | 18/19 | Coastal Norway | 1 | 1 | 0.00E+00 | 1 | 0.00E+00 | | |

Table_S3

| | | | | | | | | | | | | | | | | | |
|---|---|---|---|---|---|---|---|---|---|---|---|---|---|---|---|---|---|
| **2018.01.061** | | 2018 | 12/4/2018 | 0.259553442 | -0.585773208 | 0 | No | 12 | 18/19 | Coastal Norway | 2.22E-16 | 2.22E-16 | 0.00E+00 | 2.22E-16 | 0.00E+00 | | |
| **2018.01.063** | | 2018 | 12/4/2018 | 287.296901 | 2.458330941 | 1 | Yes | 12 | 18/19 | Coastal Norway | 1 | 1 | 0.00E+00 | 1 | 0.00E+00 | | |
| **2019.01.001** | | 2019 | 1/8/2019 | 196.0461137 | 2.292358237 | 1 | Yes | 1 | 18/19 | Coastal Norway | 1 | 1 | 0.00E+00 | 1 | 0.00E+00 | | |
| **2019.01.002** | | 2019 | 1/8/2019 | 97.59533626 | 1.989429065 | 1 | Yes | 1 | 18/19 | Coastal Norway | 1 | 1 | 0.00E+00 | 1 | 0.00E+00 | | |
| **2019.01.003** | | 2019 | 1/8/2019 | 0.317329908 | -0.498488995 | 0 | No | 1 | 18/19 | Coastal Norway | 2.22E-16 | 2.22E-16 | 0.00E+00 | 2.22E-16 | 0.00E+00 | | |
| **2019.01.004** | | 2019 | 1/9/2019 | 0.844003815 | -0.07365559 | 0 | No | 1 | 18/19 | Coastal Norway | 2.22E-16 | 2.22E-16 | 0.00E+00 | 2.22E-16 | 0.00E+00 | | |
| **2019.01.005** | | 2019 | 1/9/2019 | 322.5546217 | 2.508603269 | 1 | Yes | 1 | 18/19 | Coastal Norway | 1 | 1 | 0.00E+00 | 1 | 0.00E+00 | | |
| **2019.01.008** | | 2019 | 1/9/2019 | 1.582404165 | 0.199317417 | 0 | No | 1 | 18/19 | Coastal Norway | 2.22E-16 | 2.22E-16 | 0.00E+00 | 2.22E-16 | 0.00E+00 | 1 | |
| **2019.01.009** | | 2019 | 1/19/2019 | 260.4045613 | 2.415648587 | 1 | Yes | 1 | 18/19 | Coastal Norway | 1 | 1 | 0.00E+00 | 1 | 0.00E+00 | | |
| **2019.01.014** | | 2019 | 1/19/2019 | 0.961315948 | -0.017133853 | 0 | No | 1 | 18/19 | Coastal Norway | 2.22E-16 | 2.22E-16 | 0.00E+00 | 2.22E-16 | 0.00E+00 | | |
| **2019.01.016** | | 2019 | 1/23/2019 | 0.503806297 | -0.297736409 | 0 | No | 1 | 18/19 | Coastal Norway | 2.22E-16 | 2.22E-16 | 0.00E+00 | 2.22E-16 | 0.00E+00 | | |
| **2019.01.017** | | 2019 | 1/24/2019 | 0.372110462 | -0.42932812 | 0 | No | 1 | 18/19 | Coastal Norway | 2.22E-16 | 2.22E-16 | 0.00E+00 | 2.22E-16 | 0.00E+00 | | |
| **2019.01.019** | Mnova18050 | 2019 | 1/24/2019 | 0.545209432 | -0.263436639 | 0 | No | 1 | 18/19 | Coastal Norway | 2.22E-16 | 2.22E-16 | 0.00E+00 | 2.22E-16 | 0.00E+00 | | |
| o135 | | 2011 | 6/6/11 | 30.78 | | | Yes | | | Coastal Norway | | | | | | | extracted in Scotland |
| o335 | | 2011 | 6/15/11 | 3.64 | | | No | | | Coastal Norway | | | | | | | extracted in Scotland |
| o335 | | 2012 | 1/21/12 | 3.42 | | | No | | | Coastal Norway | | | | | | 1 | extracted in Scotland |
| o435 | | 2012 | 1/21/12 | 3.27 | | | No | | | Coastal Norway | | | | | | 1 | extracted in Scotland |
| o335 | | 2012 | 6/13/12 | 3.49 | | | No | | | Coastal Norway | | | | | | | extracted in Scotland |
| o435 | | 2012 | 6/13/12 | 3.23 | | | No | | | Coastal Norway | | | | | | | extracted in Scotland |
| o735 | | 2012 | 6/18/12 | 3.42 | | | No | | | Coastal Norway | | | | | | | extracted in Scotland |
| o635 | | 2012 | 6/18/12 | 15.67 | | | Yes | | | Coastal Norway | | | | | | | extracted in Scotland |
| o935 | | 2012 | 6/25/12 | 3.63 | | | No | | | Coastal Norway | | | | | | | extracted in Scotland |
| 1035 | | 2012 | 6/25/12 | 6.67 | | | Uncertain | | | Coastal Norway | | | | | | | extracted in Scotland |
| 1235 | | 2012 | 6/29/12 | 3.49 | | | No | | | Coastal Norway | | | | | | | extracted in Scotland |
| 1135 | | 2012 | 6/29/12 | 4.93 | | | No | | | Coastal Norway | | | | | | | extracted in Scotland |
| o135 | | 2013 | 11/29/13 | 3.53 | | | No | | | Coastal Norway | | | | | | | extracted in Scotland |
| o735 | | 2013 | 12/6/13 | 10.46 | | | Yes | | | Coastal Norway | | | | | | | extracted in Scotland |
| 1035 | | 2013 | 12/7/13 | 12.31 | | | Yes | | | Coastal Norway | | | | | | | extracted in Scotland |
| o135 | | 2014 | 1/26/14 | 3.6 | | | No | | | Coastal Norway | | | | | | | extracted in Scotland |
| o235 | | 2014 | 1/26/14 | 3.52 | | | No | | | Coastal Norway | | | | | | | extracted in Scotland |
| o635 | | 2014 | 1/31/14 | 3.47 | | | No | | | Coastal Norway | | | | | | | extracted in Scotland |
| o735 | | 2014 | 1/31/14 | 3.39 | | | No | | | Coastal Norway | | | | | | | extracted in Scotland |
| LK-Mn-04 | | 2016 | 1/20/16 | 3.42 | | | No | | | Coastal Norway | | | | | | | extracted in Scotland |
| CH-Mn-05 | | 2016 | 1/21/16 | 2.56 | | | No | | | Coastal Norway | | | | | | | extracted in Scotland |
| LK-Mn-07 | | 2016 | 1/24/16 | 2.6 | | | No | | | Coastal Norway | | | | | | | extracted in Scotland |
| LK-Mn-09 | | 2016 | 1/25/16 | 2.65 | | | No | | | Coastal Norway | | | | | | | extracted in Scotland |
| CH-Mn-10 | | 2016 | 1/26/16 | 2.72 | | | No | | | Coastal Norway | | | | | | | extracted in Scotland |
| MW-Mn-05 | | 2017 | 1/22/17 | 8.64 | | | Uncertain | | | Coastal Norway | | | | | | | extracted in Scotland |
| LK-Mn-04 | | 2017 | 1/22/17 | 30.76 | | | Yes | | | Coastal Norway | | | | | | | extracted in Scotland |
| LK-Mn-03 | | 2017 | 1/22/17 | 2.84 | | | No | | | Coastal Norway | | | | | | | extracted in Scotland |
| MB-Mn-07 | | 2017 | 1/26/17 | 23.77 | | | Yes | | | Coastal Norway | | | | | | | extracted in Scotland |
| 1035 | | 2013 | 12.07.2013 | 39.48 | 1.64 | 1 | Yes | 7 | 2013 | Coastal Norway | 1 | 1 | 0 | 1 | 0 | | sample from Scotland, re-extracted in Tromsø |
| o135 | | 2011 | 06.06.2011 | 240.22 | 2.38 | 1 | Yes | 6 | 2011 | Coastal Norway | 1 | 1 | 0 | 1 | 0 | | sample from Scotland, re-extracted in Tromsø |
| o135 | | 2011 | 06.06.2011 | 124.98 | 2.1 | 1 | Yes | 6 | 2011 | Coastal Norway | 1 | 1 | 0 | 1 | 0 | | sample from Scotland, re-extracted in Tromsø |
| LK-Mn-04 | | 2017 | 22.01.2017 | 111.66 | 2.05 | 1 | Yes | 1 | 2016/17 | Coastal Norway | 1 | 1 | 0 | 1 | 0 | | sample from Scotland, re-extracted in Tromsø |
| LK-Mn-04 | | 2017 | 22.01.2017 | 53.78 | 1.73 | 1 | Yes | 1 | 2016/17 | Coastal Norway | 1 | 1 | 0 | 1 | 0 | | sample from Scotland, re-extracted in Tromsø |
| MB-Mn-07 | | 2017 | 26.01.2017 | 162.44 | 2.21 | 1 | Yes | 1 | 2016/17 | Coastal Norway | 1 | 1 | 0 | 1 | 0 | | sample from Scotland, re-extracted in Tromsø |
| MB-Mn-07 | | 2017 | 26.01.2017 | 188.58 | 2.28 | 1 | Yes | 1 | 2016/17 | Coastal Norway | 1 | 1 | 0 | 1 | 0 | | sample from Scotland, re-extracted in Tromsø |
| o135 | | 2014 | 26.01.2014 | 4.32 | 0.64 | 0 | No | 1 | 2013/14 | Coastal Norway | 2.22E-16 | 1.33E-14 | 0 | 2.22E-16 | 0.00E+00 | | sample from Scotland, re-extracted in Tromsø |
| o335 | | 2012 | 13.06.2012 | 4.97 | 0.7 | 0 | No | 6 | 2012 | Coastal Norway | 2.22E-16 | 5.03E-12 | 0 | 2.22E-16 | 0.00E+00 | | sample from Scotland, re-extracted in Tromsø |
| o735 | | 2012 | 18.06.2012 | 2.42 | 0.38 | 0 | No | 6 | 2012 | Coastal Norway | 2.22E-16 | 2.22E-16 | 0 | 2.22E-16 | 0.00E+00 | | sample from Scotland, re-extracted in Tromsø |
| o635 | | 2012 | 18.06.2012 | 94.09 | 1.97 | 1 | Yes | 6 | 2012 | Coastal Norway | 1 | 1 | 0 | 1 | 0.00E+00 | | sample from Scotland, re-extracted in Tromsø |
| o935 | | 2012 | 25.06.2012 | 1.89 | 0.28 | 0 | No | 6 | 2012 | Coastal Norway | 2.22E-16 | 2.22E-16 | 0 | 2.22E-16 | 0.00E+00 | | sample from Scotland, re-extracted in Tromsø |
| 1235 | | 2012 | 29.06.2012 | 16.96 | 1.23 | NA | Undeterminec | 6 | 2012 | Coastal Norway | 0.999991739 | 1 | 8.26E-06 | 0.665298176 | 0.334693563 | | sample from Scotland, re-extracted in Tromsø |
| o235 | | 2014 | 26.01.2014 | 1.67 | 0.22 | 0 | No | 1 | 2013/14 | Coastal Norway | 2.22E-16 | 2.22E-16 | 0 | 2.22E-16 | 0.00E+00 | | sample from Scotland, re-extracted in Tromsø |

### Table_S3

| | | | | | | | | | | | | | | | | | |
|---|---|---|---|---|---|---|---|---|---|---|---|---|---|---|---|---|---|
| o235 | | 2014 | 26.01.2014 | 29.98 | 1.48 | 1 | Yes | 1 | 2013/14 | Coastal Norway | 1 | 1 | 0 | 1 | 0.00E+00 | | sample from Scotland, re-extracted in Tromsø |
| LK-Mn-09 | | 2016 | 25.01.2016 | 5.17 | 0.71 | 0 | No | 1 | 2015/16 | Coastal Norway | 2.22E-16 | 1.39E-11 | 0 | 2.22E-16 | 0.00E+00 | | sample from Scotland, re-extracted in Tromsø |

**Text S1 Comparison of extraction methods.** Details on the difference between the two different extractions used for progesterone analysis.

We pooled results from two slightly different extraction and quantification methods, so we confirmed that the results were comparable by repeating extraction and pregnancy assignment of a subset of samples (N = 13) using both methods. While progesterone levels differed, the same pregnancy status was assigned for all 13 of these individuals, and therefore we included all samples in the pregnancy rate analyses. We repeated the extraction and measurements for a subset of the blubber samples, in which case we report the averaged resulting progesterone level, and ran multiple samples at several dilutions.

The mean progesterone concentration for pregnant individuals was 224.4 ± 173.6 nanogram per gram of blubber (ng $g^{-1}$ P4) (NO 221.7 ± 182.2; BS 194.7 ± 7), ranging between 79.6 (194.7 BS) and 872.5 (303.8 BS). Using the other method (marked as "Scotland" in Table S3), pregnant female concentrations were 20.6 on average and ranged from 10.5-30.8. The mean concentration for non-pregnant females was 0.7 ± 0.3 ng.g-1 (NO 0.7 ± 0.3; BS 0.52 ± 0.3), ranging between 0.3 and 6.8 (0.2 - 1.1 BS); and 3.3 on average with a range from 2.6-4.9.

**Text S2 Relatedness methods.** Detailed methods for within-season re-sampling assessment using $r_{xy}$

To look for potential re-sampling individuals within sampling season, 107 genomes were sequenced at low coverage using an Illumina HiSeq4000 platform (Novogene, Hong Kong). The 1,781,057,402 paired-end reads obtained were mapped to the available humpback whale reference genome (GenBank assembly accession: GCA_004329385.1). NGSRelate v2 which allows to calculate pairwise relatedness (r) between two individuals x and y using identity by descent have been used. As input for NGSrelate, genotype likelihoods (GL) of the dataset were calculated from the mapped bam files using ANGSD v0.935-53-gf475f10. The following filtering options were used: minimum mapping and base quality of 15 (-minmapQ and -minQ 15); calculate genotype likelihoods using model from SAMtools (-GL 1); output binary genotype likelihoods (-doGlf 3); infer major and minor alleles using GL (-doMajorMinor 1); calculate per site frequencies using a fixed major and minor allele (-doMaf 1); only include SNPs with a p-value less than $1 \times 10{-}6$ (-SNP_pval 1e-6), and a minimum minor allele frequency of 0.01 (-minmaf 0.01). The number of sites used between pairwise comparisons to calculate relatedness ($r_{xy}$) ranged from 14829 to 28014.

**Table S4: Pairwise coefficients of relatedness matrix** Resulting pairwise relatedness coefficients for a subset of individuals.

## 2015/2016

| | **M16004** | **M16006** | **M16007** | **M16008** | **M16009** | **M16010** | **M16011** | **M16013** | **M16014** | **M16015** | **M16016** | **M16017** | **M16018** | **M16019** | **M16020** | **M16021** | **M16022** | **M16023** |
|---|---|---|---|---|---|---|---|---|---|---|---|---|---|---|---|---|---|---|
| **M16003** | 0.413 | 0.311 | 0.263 | 0.442 | 0.382 | 0.423 | 0.447 | 0.374 | 0.434 | 0.433 | 0.423 | 0.326 | 0.436 | 0.398 | 0.456 | 0.458 | 0.436 | 0.425 |
| **M16004** | | 0.331 | 0.292 | 0.383 | 0.367 | 0.361 | 0.328 | 0.389 | 0.342 | 0.365 | 0.326 | 0.318 | 0.360 | 0.331 | 0.389 | 0.383 | 0.360 | 0.345 |
| **M16006** | | | 0.375 | 0.338 | 0.331 | 0.344 | 0.313 | 0.311 | 0.292 | 0.300 | 0.295 | 0.363 | 0.291 | 0.293 | 0.278 | 0.303 | 0.270 | 0.284 |
| **M16007** | | | | 0.296 | 0.312 | 0.312 | 0.194 | 0.191 | 0.199 | 0.252 | 0.225 | 0.364 | 0.211 | 0.241 | 0.247 | 0.229 | 0.221 | 0.184 |
| **M16008** | | | | | 0.410 | 0.397 | 0.413 | 0.336 | 0.399 | 0.414 | 0.407 | 0.341 | 0.404 | 0.361 | 0.390 | 0.418 | 0.401 | 0.402 |
| **M16009** | | | | | | 0.417 | 0.385 | 0.348 | 0.380 | 0.386 | 0.390 | 0.349 | 0.379 | 0.364 | 0.363 | 0.367 | 0.355 | 0.373 |
| **M16010** | | | | | | | 0.431 | 0.346 | 0.388 | 0.376 | 0.392 | 0.360 | 0.408 | 0.375 | 0.384 | 0.395 | 0.377 | 0.383 |
| **M16011** | | | | | | | | 0.393 | 0.470 | 0.423 | 0.496 | 0.295 | 0.472 | 0.419 | 0.463 | 0.493 | 0.451 | 0.464 |
| **M16013** | | | | | | | | | 0.401 | 0.364 | 0.385 | 0.282 | 0.393 | 0.341 | 0.364 | 0.398 | 0.371 | 0.391 |
| **M16014** | | | | | | | | | | 0.458 | 0.500 | 0.311 | 0.441 | 0.442 | 0.443 | 0.463 | 0.444 | 0.458 |
| **M16015** | | | | | | | | | | | 0.468 | 0.331 | 0.397 | 0.434 | 0.447 | 0.455 | 0.444 | 0.424 |
| **M16016** | | | | | | | | | | | | 0.287 | 0.444 | 0.423 | 0.457 | 0.460 | 0.459 | 0.468 |
| **M16017** | | | | | | | | | | | | | 0.265 | 0.292 | 0.279 | 0.285 | 0.269 | 0.288 |
| **M16018** | | | | | | | | | | | | | | 0.429 | 0.460 | 0.447 | 0.459 | 0.460 |
| **M16019** | | | | | | | | | | | | | | | 0.455 | 0.420 | 0.430 | 0.427 |
| **M16020** | | | | | | | | | | | | | | | | 0.464 | 0.497 | 0.464 |
| **M16021** | | | | | | | | | | | | | | | | | 0.482 | 0.450 |
| **M16022** | | | | | | | | | | | | | | | | | | 0.544 |

## 2016/2017

| | **M16002** | **M16024** | **M16025** | **M16026** | **M16027** | **M16028** | **M16029** | **M17007** | **M17008** | **M17010** | **M17011** | **M17012** | **M17013** | **M17014** | **M17015** |
|---|---|---|---|---|---|---|---|---|---|---|---|---|---|---|---|
| **M16001** | 0.431 | 0.278 | 0.385 | 0.407 | 0.379 | 0.372 | 0.331 | 0.380 | 0.413 | 0.416 | 0.332 | 0.374 | 0.304 | 0.359 | 0.381 |
| **M16002** | | 0.232 | 0.434 | 0.445 | 0.454 | 0.374 | 0.305 | 0.422 | 0.423 | 0.437 | 0.301 | 0.364 | 0.270 | 0.376 | 0.436 |
| **M16024** | | | 0.161 | 0.156 | 0.167 | 0.336 | 0.391 | 0.148 | 0.177 | 0.149 | 0.391 | 0.217 | 0.398 | 0.138 | 0.149 |
| **M16025** | | | | 0.471 | 0.487 | 0.303 | 0.257 | 0.489 | 0.476 | 0.499 | 0.210 | 0.368 | 0.179 | 0.437 | 0.525 |
| **M16026** | | | | | 0.435 | 0.314 | 0.256 | 0.469 | 0.482 | 0.461 | 0.236 | 0.372 | 0.194 | 0.409 | 0.478 |
| **M16027** | | | | | | 0.311 | 0.254 | 0.461 | 0.490 | 0.442 | 0.217 | 0.382 | 0.208 | 0.430 | 0.478 |
| **M16028** | | | | | | | 0.362 | 0.302 | 0.318 | 0.305 | 0.367 | 0.326 | 0.332 | 0.266 | 0.301 |
| **M16029** | | | | | | | | 0.253 | 0.274 | 0.268 | 0.364 | 0.306 | 0.387 | 0.251 | 0.262 |

0.450 0.439 0.265 0.395 0.234 0.454 0.474 **M17007**

0.449 0.250 0.372 0.216 0.445 0.486 **M17008**

0.241 0.383 0.191 0.415 0.461 **M17010**

0.274 0.386 0.226 0.226 **M17011**

0.283 0.369 0.400 **M17012**

0.219 0.196 **M17013**

0.481 **M17014**

## 2017/2018

**M17002 M17004 M17005 M17006 M17009**

0.466 0.469 0.485 0.465 0.442 **M17001**

0.452 0.489 0.475 0.426 **M17002**

0.422 0.418 0.442 **M17004**

0.475 0.388 **M17005**

0.419 **M17006**

## 2018/2019

**M18033 M18034 M18035 M18036 M18037 M18038 M18039 M18040 M18041 M18042 M18043 M18044 M18045 M18046 M18047 M18048 M18049 M18051 M18052 M18053 M18054 M18055 M18056 M18058 M18061 M18065 M18066 M18067 M18068 M18069 M18070 M18071 M18074 M18075 M18076 M18077**

**M18078 M18079 M18080 M18081 M18082** 0.221 0.484 0.485 0.504 0.489 0.386 0.487 0.532 0.474 0.475 0.463 0.332 0.398 0.477 0.497 0.210 0.467 0.490 0.217 0.259 0.487 0.243 0.125 0.140 0.235 0.544 0.317 0.398 0.389 0.352 0.404 0.477 0.322 0.278 0.495 0.434 0.474 0.428 0.468 0.394 0.463 **M18001** 0.253 0.243 0.261

0.250 0.269 0.234 0.244 0.233 0.257 0.241 0.340 0.294 0.178 0.247 0.400 0.257 0.253 0.402 0.413 0.217 0.384 0.368 0.356 0.379 0.221 0.356 0.322 0.306 0.403 0.317 0.233 0.382 0.434 0.290 0.308 0.290 0.299 0.266 0.315 0.252 **M18033** 0.635 0.505 0.491 0.384 0.463 0.489 0.466 0.460 0.454 0.373 0.424 0.449 0.499 0.244 0.450

0.476 0.243 0.264 0.442 0.264 0.136 0.157 0.250 0.505 0.341 0.424 0.404 0.340 0.419 0.440 0.321 0.285 0.479 0.413 0.443 0.415 0.465 0.431 0.495 **M18034** 0.509 0.493 0.393 0.470 0.505 0.463 0.457 0.452 0.361 0.421 0.473 0.520 0.234 0.451 0.485 0.240 0.256 0.452 0.248 0.126 0.157 0.238 0.524 0.312 0.426 0.387 0.342 0.420

0.455 0.318 0.292 0.484 0.394 0.449 0.413 0.465 0.427 0.504 **M18035** 0.477 0.376 0.473 0.525 0.476 0.506 0.467 0.372 0.421 0.458 0.523 0.255 0.475 0.514 0.256 0.264 0.457 0.256 0.132 0.142 0.254 0.527 0.319 0.429 0.413 0.371 0.423 0.476 0.345 0.296 0.480 0.407 0.451 0.438 0.457 0.409 0.473 **M18036** 0.370 0.484 0.502

0.464 0.449 0.471 0.351 0.457 0.461 0.498 0.212 0.448 0.493 0.232 0.258 0.443 0.247 0.152 0.140 0.233 0.513 0.324 0.419 0.391 0.352 0.388 0.481 0.326 0.273 0.481 0.449 0.447 0.440 0.463 0.406 0.475 **M18037** 0.393 0.393 0.381 0.391 0.382 0.341 0.374 0.372 0.359 0.273 0.376 0.410 0.272 0.298 0.360 0.301 0.202 0.218 0.284

0.390 0.316 0.353 0.367 0.310 0.329 0.374 0.335 0.316 0.367 0.361 0.368 0.350 0.347 0.349 0.383 **M18038** 0.478 0.493 0.500 0.497 0.379 0.422 0.467 0.499 0.204 0.463 0.481 0.213 0.261 0.464 0.284 0.127 0.126 0.237 0.473 0.343 0.399 0.407 0.322 0.381 0.489 0.326 0.270 0.482 0.391 0.430 0.424 0.431 0.384 0.460 **M18039**

0.496 0.484 0.474 0.356 0.423 0.465 0.514 0.219 0.484 0.527 0.214 0.262 0.455 0.251 0.147 0.162 0.256 0.515 0.321 0.420 0.410 0.352 0.421 0.496 0.351 0.285 0.499 0.431 0.454 0.422 0.480 0.422 0.498 **M18040** 0.493 0.482 0.372 0.409 0.506 0.488 0.218 0.497 0.480 0.219 0.250 0.473 0.265 0.131 0.125 0.239 0.478 0.321 0.412

0.389 0.336 0.395 0.505 0.341 0.287 0.500 0.411 0.435 0.410 0.454 0.395 0.450 **M18041** 0.468 0.388 0.425 0.450 0.481 0.231 0.473 0.486 0.233 0.245 0.464 0.275 0.150 0.136 0.233 0.467 0.352 0.411 0.390 0.356 0.401 0.477 0.336 0.308 0.462 0.395 0.442 0.413 0.439 0.400 0.459 **M18042** 0.360 0.423 0.454 0.488 0.204 0.475

0.495 0.215 0.238 0.461 0.263 0.123 0.127 0.235 0.469 0.304 0.387 0.369 0.334 0.381 0.481 0.317 0.276 0.450 0.396 0.423 0.432 0.441 0.399 0.465 **M18043**

0.378 0.283 0.368 0.330 0.367 0.360 0.332 0.338 0.316 0.354 0.286 0.260 0.326 0.327 0.394 0.387 0.391 0.326 0.404 0.356 0.327 0.347 0.366 0.352 0.387 0.384 0.338 0.391 0.370 **M18044**

0.352 0.447 0.259 0.413 0.444 0.273 0.286 0.365 0.300 0.201 0.192 0.277 0.405 0.356 0.419 0.401 0.349 0.399 0.398 0.334 0.301 0.389 0.412 0.413 0.405 0.398 0.411 0.402 **M18045**

0.504 0.177 0.466 0.463 0.168 0.216 0.478 0.190 0.096 0.109 0.197 0.521 0.256 0.351 0.338 0.311 0.341 0.485 0.280 0.229 0.476 0.362 0.385 0.355 0.416 0.339 0.441 **M18046**

0.215 0.480 0.489 0.228 0.242 0.477 0.245 0.103 0.122 0.211 0.539 0.327 0.421 0.371 0.369 0.414 0.473 0.346 0.278 0.481 0.416 0.437 0.431 0.495 0.427 0.495 **M18047**

0.213 0.207 0.411 0.400 0.194 0.381 0.382 0.414 0.394 0.211 0.344 0.294 0.291 0.313 0.285 0.190 0.327 0.398 0.264 0.265 0.269 0.266 0.206 0.300 0.230 **M18048**

0.544 0.231 0.268 0.464 0.269 0.151 0.141 0.249 0.472 0.344 0.421 0.393 0.372 0.405 0.482 0.350 0.304 0.489 0.429 0.450 0.445 0.468 0.405 0.456 **M18049**

0.225 0.263 0.451 0.250 0.128 0.128 0.256 0.508 0.329 0.429 0.417 0.359 0.417 0.468 0.326 0.283 0.474 0.436 0.474 0.443 0.465 0.415 0.480 **M18051**

0.401 0.216 0.388 0.361 0.380 0.390 0.208 0.327 0.302 0.284 0.351 0.282 0.215 0.343 0.401 0.245 0.276 0.263 0.282 0.247 0.294 0.259 **M18052**

0.246 0.406 0.357 0.380 0.398 0.245 0.320 0.317 0.313 0.364 0.317 0.248 0.350 0.413 0.284 0.305 0.298 0.296 0.263 0.320 0.275 **M18053**

0.254 0.120 0.116 0.209 0.508 0.310 0.369 0.340 0.337 0.366 0.500 0.327 0.281 0.415 0.402 0.433 0.379 0.458 0.391 0.455 **M18054**

0.370 0.356 0.355 0.224 0.347 0.315 0.307 0.328 0.306 0.273 0.321 0.382 0.259 0.289 0.285 0.282 0.269 0.320 0.252 **M18055**

0.408 0.361 0.097 0.305 0.222 0.224 0.255 0.212 0.193 0.265 0.337 0.169 0.198 0.175 0.189 0.135 0.213 0.142 **M18056**

0.362 0.133 0.290 0.197 0.201 0.266 0.211 0.145 0.281 0.356 0.203 0.219 0.168 0.214 0.147 0.225 0.176 **M18058**

0.216 0.319 0.315 0.384 0.310 0.311 0.219 0.302 0.380 0.252 0.260 0.282 0.300 0.242 0.283 0.241 **M18061**

0.302 0.390 0.374 0.343 0.386 0.467 0.334 0.256 0.477 0.426 0.465 0.397 0.499 0.398 0.485 **M18065**

0.400 0.368 0.325 0.381 0.319 0.345 0.368 0.335 0.355 0.377 0.368 0.326 0.378 0.334 **M18066**

0.429 0.362 0.418 0.407 0.350 0.334 0.433 0.418 0.441 0.427 0.416 0.428 0.410 **M18067**

0.328 0.420 0.379 0.314 0.310 0.412 0.370 0.427 0.424 0.371 0.397 0.396 **M18068**

0.351 0.330 0.496 0.398 0.369 0.358 0.359 0.378 0.346 0.354 0.362 **M18069**

0.382 0.314 0.352 0.417 0.418 0.427 0.431 0.414 0.425 0.420 **M18070**

0.329 0.291 0.449 0.396 0.416 0.389 0.455 0.380 0.447 **M18071**

0.398 0.348 0.357 0.354 0.337 0.350 0.346 0.334 **M18074**

0.318 0.325 0.323 0.327 0.298 0.346 0.299 **M18075**

0.436 0.445 0.419 0.459 0.439 0.484 **M18076**

0.442 0.395 0.449 0.431 0.421 **M18077**

0.430 0.437 0.415 0.440 **M18078**

0.409 0.424 0.425 **M18079**

0.428 0.474 **M18080**

0.550 **M18081**

## S2009

**M18003 M18004 M18005 M18006 M18007 M18008 M18009 M18010 M18011 M18012 M18013 M18014 M18015 M18016 M18018 M18019 M18028 M18029**
**M18030 M18031** 0.202 0.464 0.452 0.464 0.433 0.467 0.459 0.461 0.488 0.463 0.492 0.473 0.514 0.475 0.463 0.273 0.480 0.458 0.464 0.435 **M18002** 0.183
0.217 0.200 0.277 0.215 0.218 0.160 0.174 0.226 0.234 0.217 0.214 0.218 0.203 0.341 0.251 0.229 0.206 0.245 **M18003** 0.462 0.443 0.415 0.479 0.459 0.465
0.449 0.462 0.455 0.442 0.458 0.474 0.495 0.263 0.461 0.481 0.452 0.450 **M18004** 0.423 0.387 0.450 0.442 0.418 0.470 0.440 0.444 0.422 0.457 0.466 0.460
0.308 0.445 0.450 0.450 0.463 **M18005** 0.399 0.467 0.454 0.397 0.466 0.465 0.450 0.472 0.441 0.440 0.431 0.316 0.450 0.430 0.456 0.415 **M18006** 0.441 0.441
0.374 0.429 0.432 0.453 0.444 0.440 0.451 0.415 0.344 0.448 0.425 0.415 0.406 **M18007** 0.492 0.414 0.479 0.502 0.482 0.486 0.484 0.479 0.488 0.303 0.472
0.473 0.469 0.458 **M18008**
0.437 0.484 0.613 0.483 0.508 0.497 0.482 0.478 0.286 0.475 0.470 0.461 0.460 **M18009**
0.409 0.444 0.421 0.455 0.411 0.411 0.447 0.215 0.420 0.436 0.412 0.409 **M18010**
0.480 0.473 0.456 0.492 0.484 0.459 0.303 0.471 0.437 0.468 0.432 **M18011**
0.481 0.503 0.499 0.470 0.481 0.284 0.463 0.464 0.464 0.465 **M18012**
0.492 0.485 0.473 0.470 0.305 0.507 0.479 0.456 0.436 **M18013**
0.471 0.453 0.453 0.287 0.471 0.465 0.476 0.416 **M18014**
0.491 0.485 0.291 0.499 0.464 0.490 0.451 **M18015**
0.465 0.318 0.468 0.440 0.448 0.425 **M18016**
0.287 0.456 0.465 0.440 0.464 **M18018**
0.303 0.303 0.287 0.314 **M18019**
0.534 0.476 0.439 **M18028**
0.461 0.464 **M18029**
0.430 **M18030**
